# Production of selenium nanoparticles occurs through an interconnected pathway of sulfur metabolism and oxidative stress response in *Pseudomonas putida* KT2440

**DOI:** 10.1101/2022.09.29.507446

**Authors:** Roberto Avendaño, Said Muñoz-Montero, Diego Rojas-Gätjens, Paola Fuentes, Sofía Vieto, Rafael Montenegro, Manuel Salvador, Rufus Frew, Juhyun Kim, Max Chavarría, Jose I. Jiménez

## Abstract

The soil bacterium *Pseudomonas putida* KT2440 has been shown to produce selenium nanoparticles aerobically from selenite; however, the molecular actors involved in this process are unknown. Here, through a combination of genetic and analytical techniques, we report the first insights into selenite metabolism in this bacterium. Our results suggest that the reduction of selenite occurs through an interconnected metabolic network involving central metabolic reactions, sulfur metabolism, and the response to oxidative stress. Genes such as *sucA*, D2HGDH and PP_3148 revealed that the 2-ketoglutarate and glutamate metabolism is important to converting selenite into selenium. On the other hand, mutants affecting the activity of sulfite reductase reduced the bacteria’s ability to transform selenite. Other genes related to sulfur metabolism (*ssuEF*, *sfnCE, sqrR*, *sqr* and *pdo2*) and stress response (*gqr*, *lsfA*, *ahpCF* and *sadI*) were also identified as involved in selenite transformations. Interestingly, suppression of genes *sqrR*, *sqr* and *pdo2* resulted in the production of selenium nanoparticles at a higher rate than the wild-type strain, which is of biotechnological interest. The data provided in this study brings us closer to understanding the metabolism of selenium in bacteria, and offers new targets for the development of biotechnological tools for the production of selenium nanoparticles.

## Introduction

Selenium (Se) is a metalloid element with multiple applications in health and industry. Nanometric selenium has shown very varied biological activities, including an anticancer, antibacterial, and protective agent for DNA against UV damage (Cremonini *et al*., 2016; Rayman, 2018; Khurana *et al*., 2019). In industry, elemental selenium is used to manufacture fertilizers, semiconductors and sensors or to be combined with other metals for electronics, images or photovoltaics (Chaudhary and Mehta, 2014). Considering the increased interest in using selenium in high-tech applications (Nayak *et al*., 2021), treatment systems should not only aim to remove dissolved selenium but also consider recovering strategies of valuable nanoparticles and move towards a circular economy. In this sense, one of the methodologies that have recently attracted attention is the use of microorganisms to carry out the bioreduction of selenium oxyanions (i.e., selenate, selenite) to elemental selenium.

Several environmental bacteria have shown the ability to perform selenate and selenite reduction. A representative example is *Thauera selenatis* (Macy *et al*., 1993) which reduces both selenate and selenite anaerobically. Other bacteria from different genera have demonstrated the ability to reduce selenite to elemental selenium aerobically. These include species of *Pseudomonas* (Avendaño *et al*., 2016), *Comamonas* (Zheng *et al*., 2014; Tan *et al*., 2018), *Enterobacter (*Song *et al*., 2017), *Azospirillum* (Tugarova *et al*., 2020), *Bacillus* (Lampis *et al*. 2014) *Shewanella* (Klonowska *et al*. 2005; Li *et al*., 2014) and *Burkholderia* (Khoei *et al*., 2017) among others.

Despite many reports have shown the potential of many bacteria for the treatment of water and soil contaminated with selenium, we still ignore many aspects of how bacteria carry out this bioreduction process. In most cases, only a few pieces of the metabolic puzzle are known and studies have been limited to only a few bacteria including *Escherichia coli*, *Salmonella typhimurium, Rhodobacter sphaeroides* and *Thauera selenatis* among others. Published works point to different selenate and selenite reduction mechanisms that include oxidative stress response (Bebien *et al*., 2001; Bébien *et al*., 2002; Kessi, 2006; Tan *et al*., 2018; Yasir *et al*., 2020) sulfur metabolism (Tan *et* al., 2018; Yasir *et al*., 2020, as well as the participation of selenate and nitrite reductases in anaerobic bacteria. (DeMoll-Decker and Macy, 1993; Schröder *et al.*, 1997; Krafft *et al*., 2000; Debieux *et al*., 2011; Butler *et al*., 2012). In *E. coli* and *Rhodospirillum rubrum* it’s been shown that glutathione is a fundamental actor in a process combining abiotic chemical reactions and enzymes (Kessi and Hanselmann, 2004).

In general terms, there is a partial knowledge of mechanisms of reduction of selenium oxyanions in bacteria; however, there are many other cases of species with potential and very attractive characteristics for the bioremediation of selenium, for which there is no mechanistic information available. One of them is *Pseudomonas putida* KT2440, a bacterium whose ability to reduce selenite to selenium was previously demonstrated (Fig. 1A) (Avendaño *et al*., 2016). This soil bacterium is particularly interesting in biotechnology and specifically in reducing processes (Nikel *et al*., 2016; Nikel and de Lorenzo, 2018). One of these characteristics is that its central carbon metabolism is adapted to produce a high reducing power (Chavarría *et al.*, 2013; Nikel *et al*., 2015; Nikel *et al.*, 2021); due to a sugar catabolism funneled through the EDEMP cycle that directs the carbon flux to NADPH-forming reactions (Nikel and Chavarría, 2015). The resulting NADPH is responsible for maintaining glutathione in its reduced form, and as a consequence, *P. putida* has high reducing power and greater resistance to oxidative stress (Chavarría *et al.*, 2013; Nikel *et al.*, 2015; Nikel *et al.*, 2021). This metabolic capacity could be used for the implementation of reduction reactions such as the transformation of oxyanions (e.g. selenite, tellurite) to their respective elemental species (Se°, Te°) (Montenegro *et al.*, 2021) (Vieto *et al.*, 2021).

**Figure 1.**
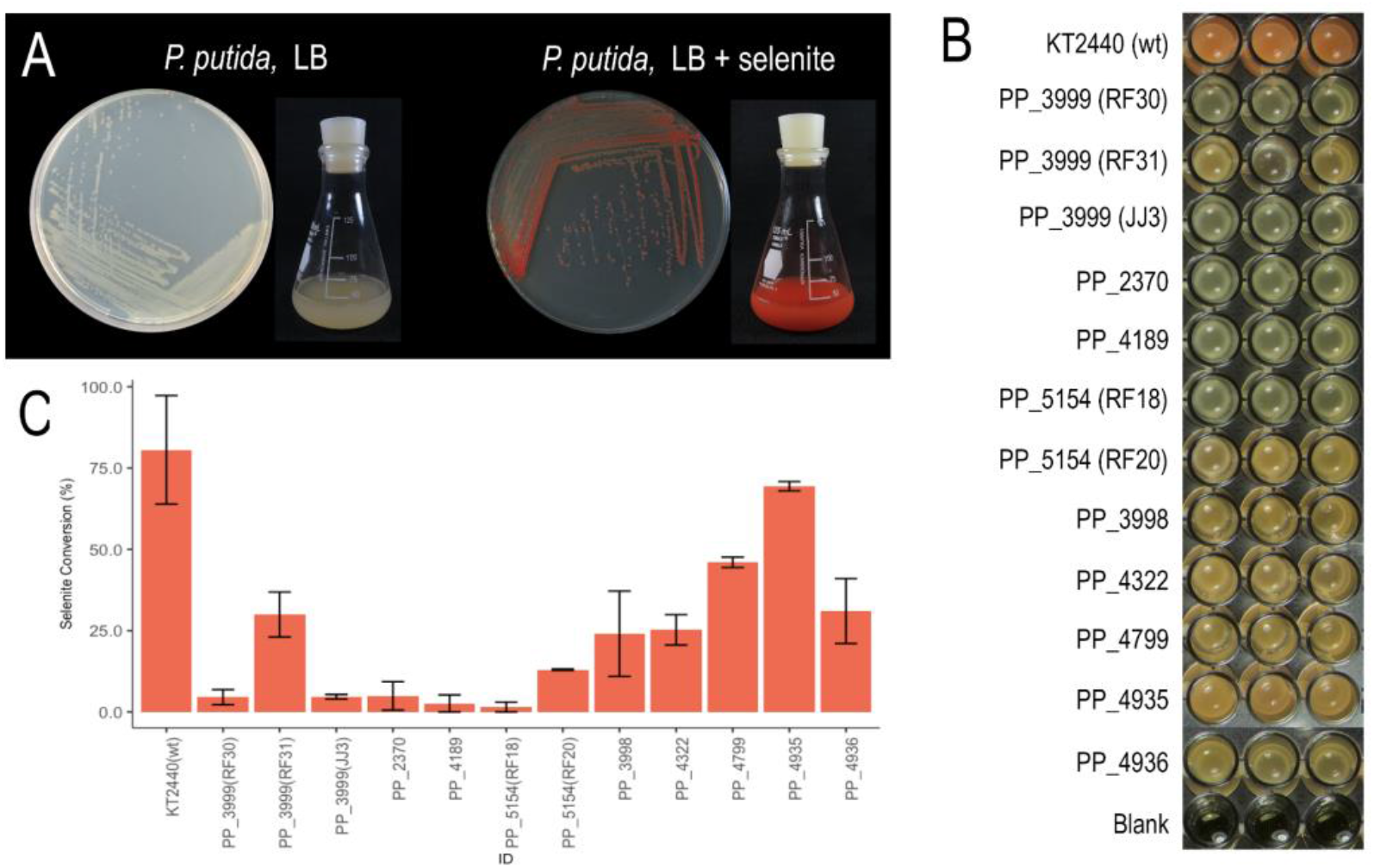
Growth of *P. putida* KT2440 and deficient strains in LB broth in the presence of selenite (1 mM). A) Photographs of *P. putida* KT2440 (wt) in both solid and liquid LB medium in the absence and presence of selenite. The photographs were taken after 24 hours of growth. B) Photographs of *P. putida* KT2440 (wt) and the twelve deficient strains grown in LB culture medium in a microplate after 24 h. C) Percentages of conversion of selenite to selenium by *P. putida* KT2440 and the twelve deficient strains in LB culture medium after 24 hours of growth. The average of three replicates is presented in the bars and the error was calculated as the standard deviation of the results.

In this work, we have identified a collection of genes involved in selenite reduction in *P. putida* KT2440 through random mutagenesis, phenotypic analyses, flame atomic absorption spectroscopy (FAAS) and RNA sequencing. Our results indicate that selenite reduction takes place by the involvement of activities related to sulfur metabolism, central metabolic reactions and the resistance to oxidative stress. We have also identified activities that compete with selenite reduction, and shown that knock outs in the corresponding genes result in faster selenium nanoparticle formation, which could lead to biotechnological applications of *P. putida* for the detoxification and valorisation of selenite waste.

## Results

### Phenotypes and growth rates of mutants involved in selenite reduction

A total of 15 mutant strains related to selenium metabolism were obtained (Table 1). Twelve strains (mutated in genes *cysG*, PP_2370, *sucA*, *D2HGDH*, *gqr*, *ccmF*, *ldcA*, *msbA* and *wzy*) had a phenotype deficient in the production of elemental selenium nanoparticles, meaning the colonies had a white to pink color during the first 24 h when cultured in LB medium with 1 mM selenite (Fig. 1B). This lack of reddish coloration denotes that these strains cannot generate elemental selenium at the same rate or yield as *P. putida* KT2440. In the remainder of the manuscript, we will refer to these strains as “deficient strains”. The rest of the mutants (*sqrR*, *sqr* and *pdo2*) turned out to be capable of producing selenium at a higher rate than the wild strain (Figs. 2A, 2B and Supp. Video S1). When observed in an LB culture with 1 mM selenite, they turned reddish noticeably faster than the wild-type strain. In the remainder of the manuscript, we will refer to these strains as “fast strains”.

**Table 1.**
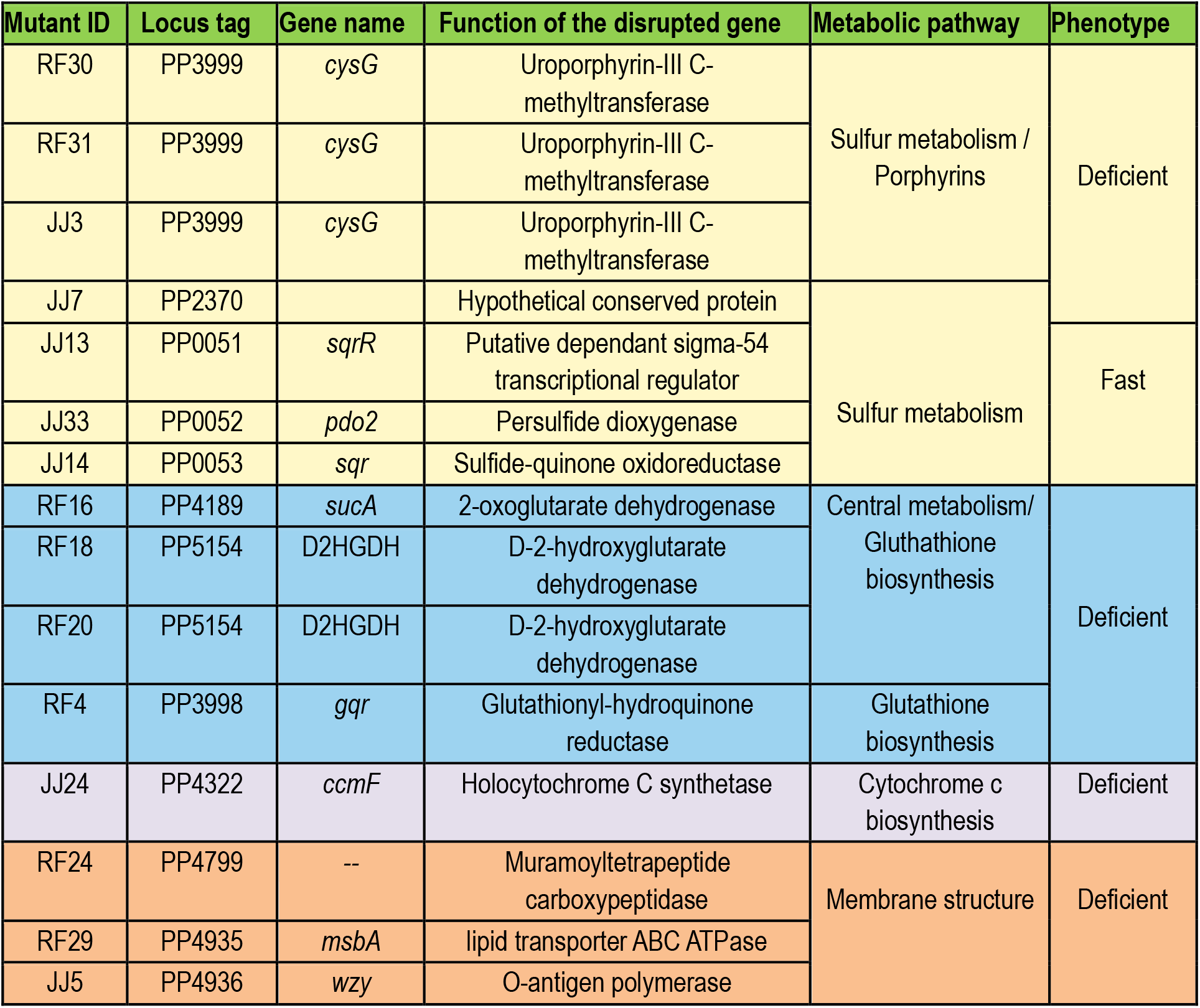
Genes mutated in *P. putida* KT2440 by insertion of kanamycin resistance cassette selected by phenotype.

**Figure 2.**
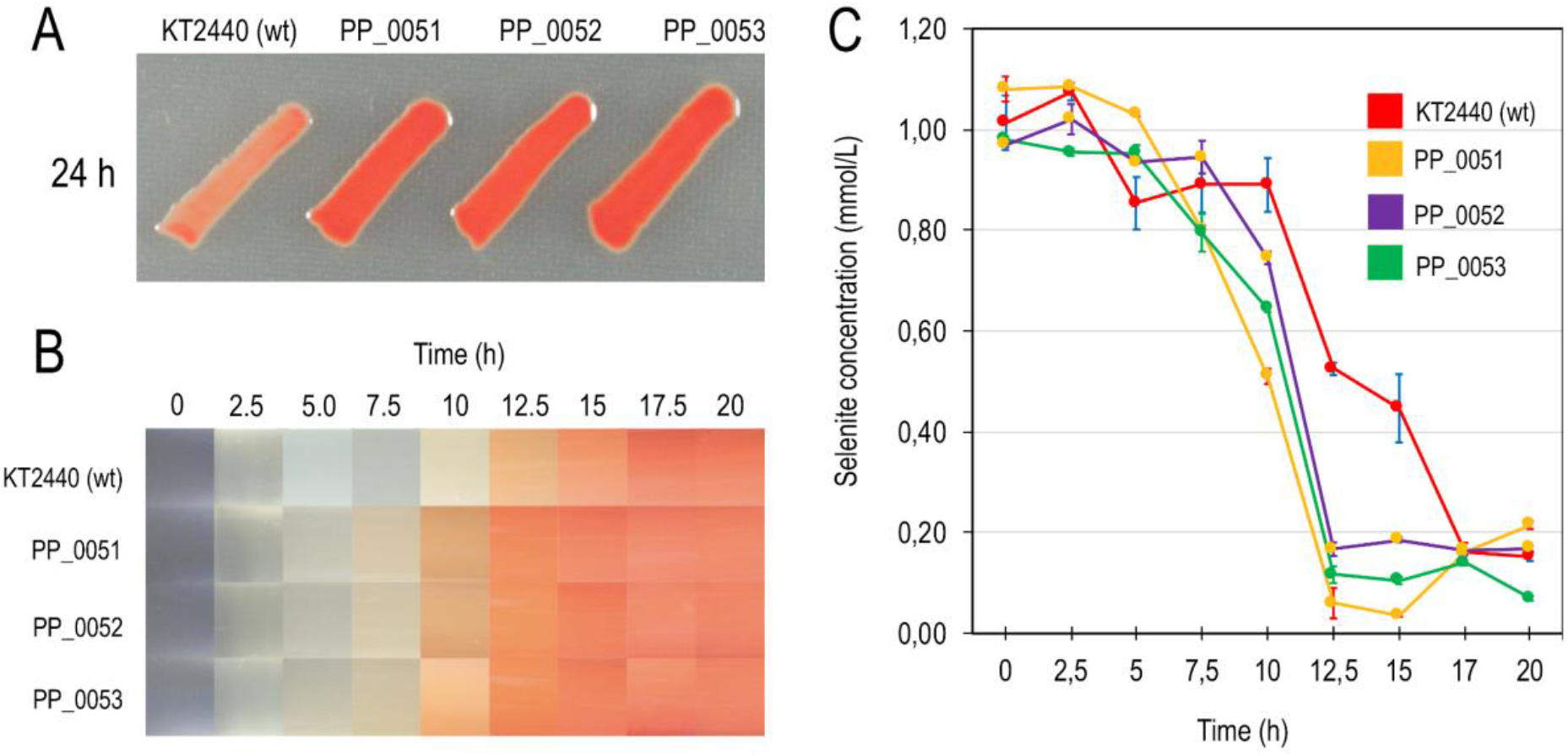
Growth of *P. putida* KT2440 and fast strains in LB broth in the presence of selenite (1 mM). A) Photographs of *P. putida* KT2440 (wt) in solid LB medium in the presence of selenite. The photographs were taken after 24 hours of growth. B) Heat map of selenite reduction in *P. putida* KT2440 and fast strains. The heat map was made using the photographs from the Supplemental Video S1 C) Kinetics of selenite reduction in *P. putida* KT2440 and fast strains measured by FAAS during 20 hours. As can be seen both in the photographs and in the reduction kinetics, the PP_0051, PP_0052 and PP_0053 strains reduce selenite more quickly.

The growth of the deficient strains in LB medium without selenite showed in all cases a growth rate similar to the wild-type strain except for the *sucA* mutant (PP_4189), which showed a slight reduction in the growth rate and the formation of biomass (Supp. Fig. S1A). This phenotype is not surprising since *sucA* encodes the enzyme 2-oxoglutarate dehydrogenase, a key enzyme in the Krebs cycle. In the presence of selenite, three strains appear to have a slower growth rate than the wild-type strain, including again the *sucA* mutant. The other two strains correspond to mutants disrupted in genes *ccmF* (encoded to holocytochrome C synthetase) and *wzy* (encoded O-antigen polymerase). In the presence of selenite, the final optical density was highly variable among strains. This phenomenon occurs because, with spectrophotometric measurements at 600 nm, elemental selenium also absorbs radiation or scatters it, so it is clear that the values measured correspond to cell growth and variable amounts of elemental selenium. As mentioned above, three mutant strains showed a very noticeable phenotype in Petri dishes and liquid cultures: these strains turn reddish faster than *P. putida* KT2440 (Figs. 2A, 2B and Supp. Video S1). These fast strains were affected by the genes *sqrR* (encoding to a putative dependent sigma-54 transcriptional regulator), *sqr* (encoding a sulfide-quinone oxidoreductase) and *pdo2* (encoding a persulfide dioxygenase). Growth curves of these mutant strains in LB medium (Supp. Fig. S2A) indicated that their growth rate is similar to the wild-type strain. In the presence of selenite, a rapid increase in optical density is observed at 600 nm due to the rapid production of selenium nanoparticles (Supp. Fig. S2B). This effect begins to be visible after 4 hours for the three strains. It is clear that the difference in the optical density observed in Supp. Fig. S2B is a product of the absorbance or dispersion of the selenium nanoparticles and not due to accelerated growth of the mutants since in the absence of selenite (Supp. Fig. S2A), no changes were observed in the growth of these strains compared to the wild-type.

The biomineralization of selenite to elemental selenium in the mutant strains was monitored by determining the amount of selenite remaining in LB cultures using Flame Atomic Absorption Spectroscopy (FAAS). For mutants deficient in selenite reduction, the presence of this oxyanion in the culture medium was determined after 24 hours of growth. As expected, all deficient strains have lower bioconversion values than the wild-type strain (Fig. 1C). However, results suggest that none of the mutated genes is essential for this process since (i) in all the cases is possible to observe some bioconversion after 24 hours (Fig. 1C) and (ii) after 48 hours of growth, all of these mutant strains manage to reach a reddish coloration almost of the same intensity as the wild-type strain (data not shown), suggesting there is more than one pathway to reduce selenite. Therefore, it is important to clarify that the term deficient does not denote a complete inability to metabolize selenite but rather denotes a limited ability to reduce selenite at the same rate as the wild-type strain.

To certify the differences in the rate of conversion of selenite between strains *sqrR*, *sqr* and *pdo2* concerning the wild-type strain, we determined in LB cultures enriched with 1 mM selenite the remaining content of oxyanion in the culture medium between 0-20 hours of growth at intervals of 2.5 h (Fig. 2C). Results showed that strains *sqrR*, *sqr* and *pdo2* start the reduction process several hours earlier than the wild-type strain; however, the yield of selenium nanoparticles is the same in all cases (wild-type and mutants) after 17.5 h (Figs. 2B, 2C and Supp. Video S1). In our previous study (Avendaño *et al.*, 2016), we concluded that the selenite reduction process in *P. putida* KT2440 begins in the middle-exponential phase; however, in *sqrR*, *sqr* and *pdo2* mutants, the reduction is activated at the beginning of the exponential phase.

### Complementation experiments in trans

In order to validate whether the observed differences in selenium formation were due to the absence of each gene individually, we ran complementation tests. For these experiments, we constructed plasmids using the pSEVA438 plasmid (Silva-Rocha *et al.*, 2013) in which genes *cysG*, *sucA*, *D2HGDH*, *gqr*, *ccmF*, *ldcA*, *msbA* and *wzy* were individually expressed under the control of a promoter inducible with 3-methylbenzoate (3MB; see Materials and Methods). These plasmids were introduced within the corresponding mutant strain, that is, each strain was complemented with its respective gene in trans. For example, for the *cysG* mutant, a plasmid pSEVa438-*cysG* was constructed and electroporated, then, induction of the gene from the plasmid was done with 3MB. As seen in Supplementary Figure S3, mutant strains with the empty vector (i.e. pSEVA438) maintain the deficient phenotype in selenite reduction (first row in the figure). However, when the mutants were complemented with the respective gene cloned in the plasmid and induced with 3MB, the selenite reduction was restored to the levels of the wild-type strain (second row in the figure). These results validate those shown in Figs. 1B and 1C and certify the involvement of these genes in selenite metabolism in *P. putida* KT2440.

### Transcriptional analysis of wild-type P. putida KT2440 in the absence and presence of selenite

We investigated the global transcriptional response to selenite of *P. putida* KT2440 preparing RNAseq libraries of cultures grown in the presence of the oxyanion.

We extracted the RNA from cultures growing on LB in the exponential phase in the absence of selenite as control or exposed to 1 mM selenite for one hour or 8 hours, the latter the condition corresponding to the maximum rate of Se° biosynthesis reported previously (Avendaño *et al.*, 2016). Libraries were prepared from two independent biological replicates in each case. We obtained an average of 17.6 million reads per sample, and after the removal of low-quality reads, 93.7% of reads were mapped to the genome of *P. putida* KT2440 (NC_002947.4)(Belda *et al.*, 2016). Reads aligned with most open reading frames except PP_1099, PP_1149 and PP_2463.

Many genes were differentially expressed in response to exposure to selenite (Fig. 3). They were classified according to their functional categories using the COG database (Fig. 4; Supp. Tables S1 and S2). We detected overexpression of 2 functional categories while 7 functional categories were underrepresented in the presence of selenite. Several functions were enriched in each condition (Fig. 5; corresponding KEGG pathways are shown in Supp Figs. S4 to S12). Exposure to selenite triggered the expression of transporters (or more broadly membrane associated proteins) and genes related to sulfur metabolism. The exposure to selenite led to the underexpression of metabolic functions involved in the metabolism of amino acids and oxidative phosphorylation. Again, the synthesis of extracellular selenium could have an impact on the membrane bound respiratory chain, preventing its correct functioning, which would affect the cellular metabolism as a whole. Moreover, the stress generated by selenite is likely to also affect metabolic processes.

**Figure 3.**
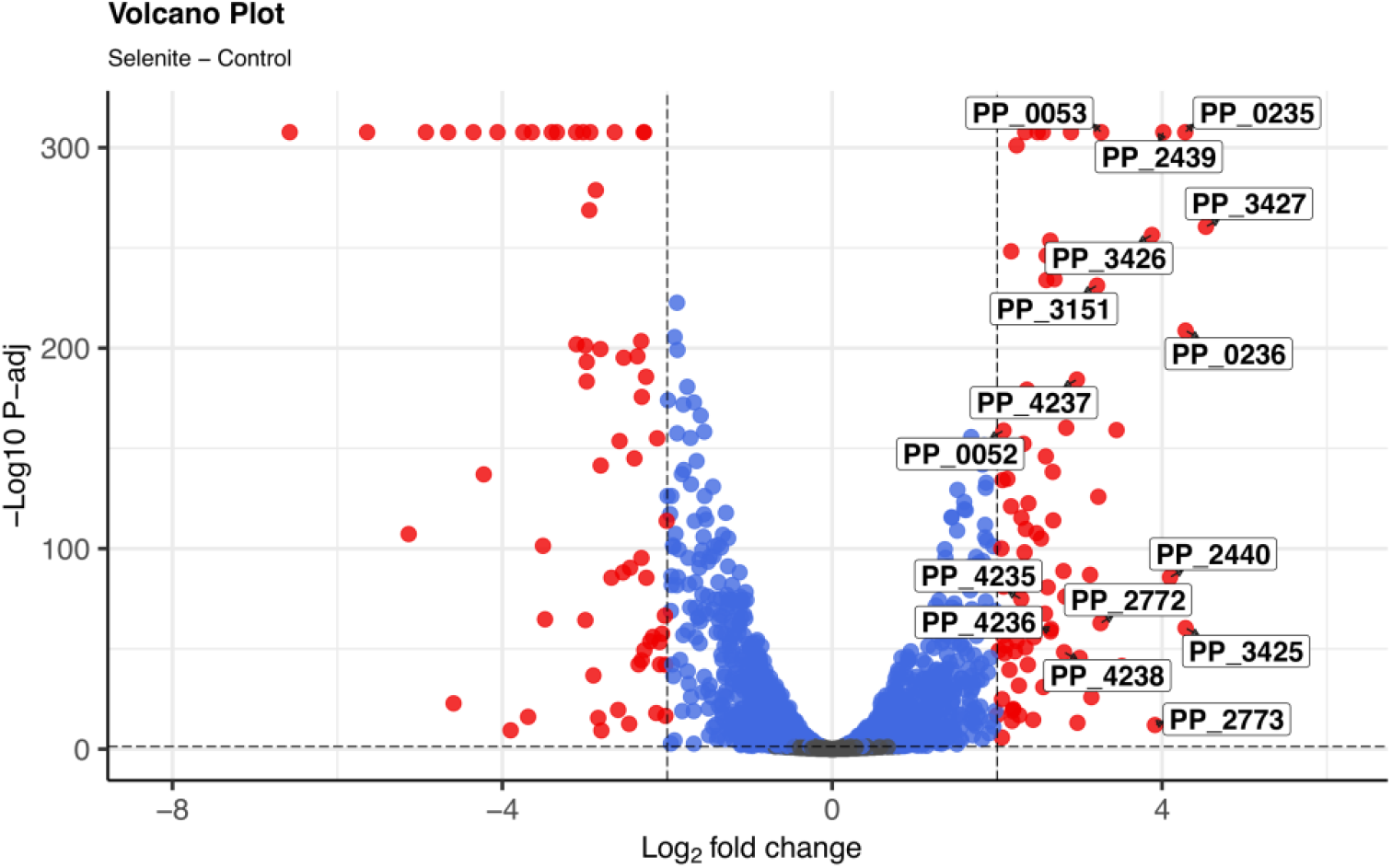
Transcriptional changes in the presence of selenite. Differential gene expression relative to control treatment is shown in the presence of selenite. Each point shows one gene. Red points show significantly differentially expressed genes (p-adjusted < 0.05 and log2 fold change > 2). Labels correspond to genes discussed in the main text.

**Figure 4.**
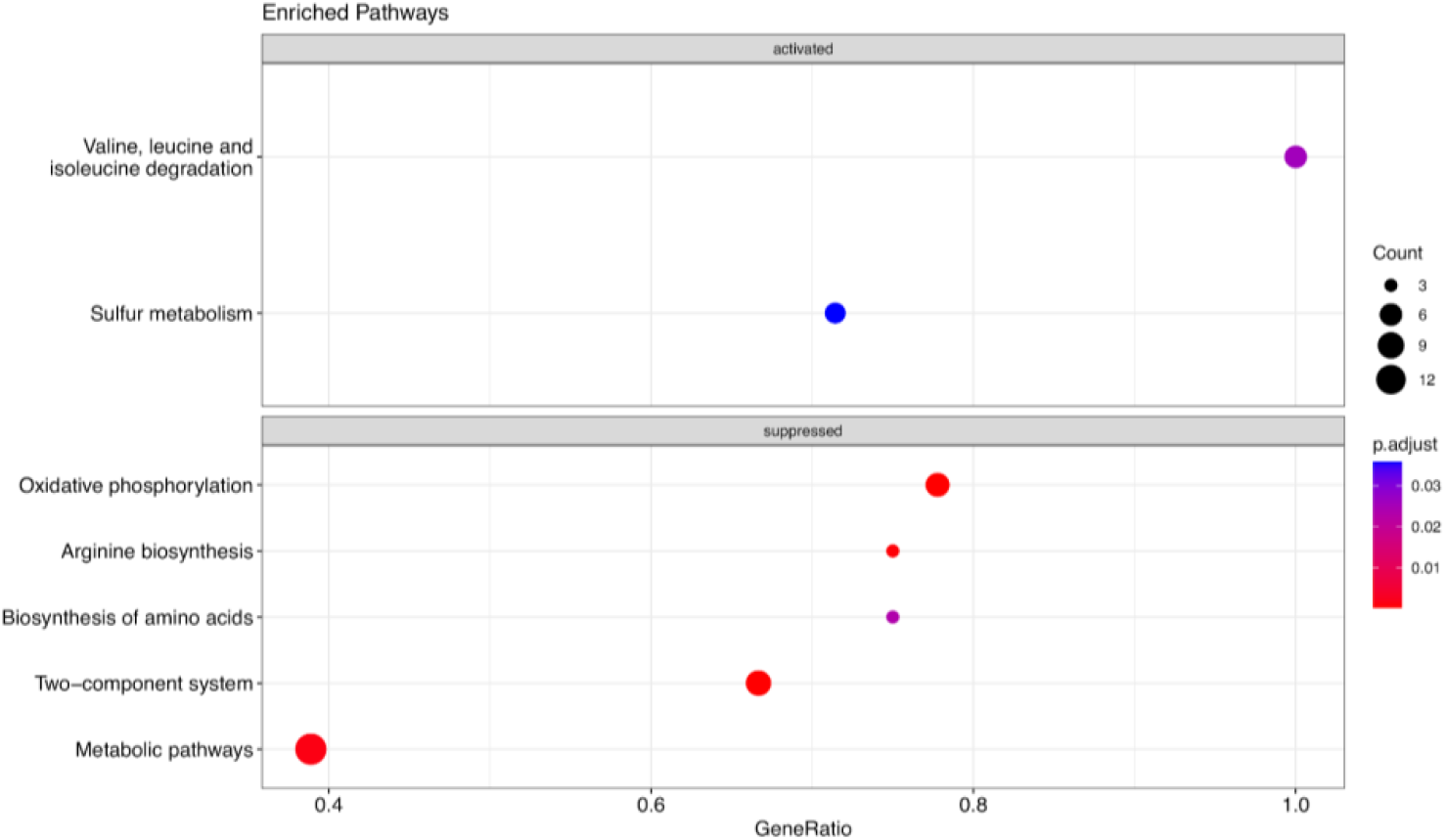
Enrichment pathways using KEGG database. Enrichment analysis was conducted comparing differentially expressed genes against the KEGG *P. putida* KT2440 database. Gene ratio corresponds to the fraction of differentially expressed genes in pathways.

**Figure 5.**
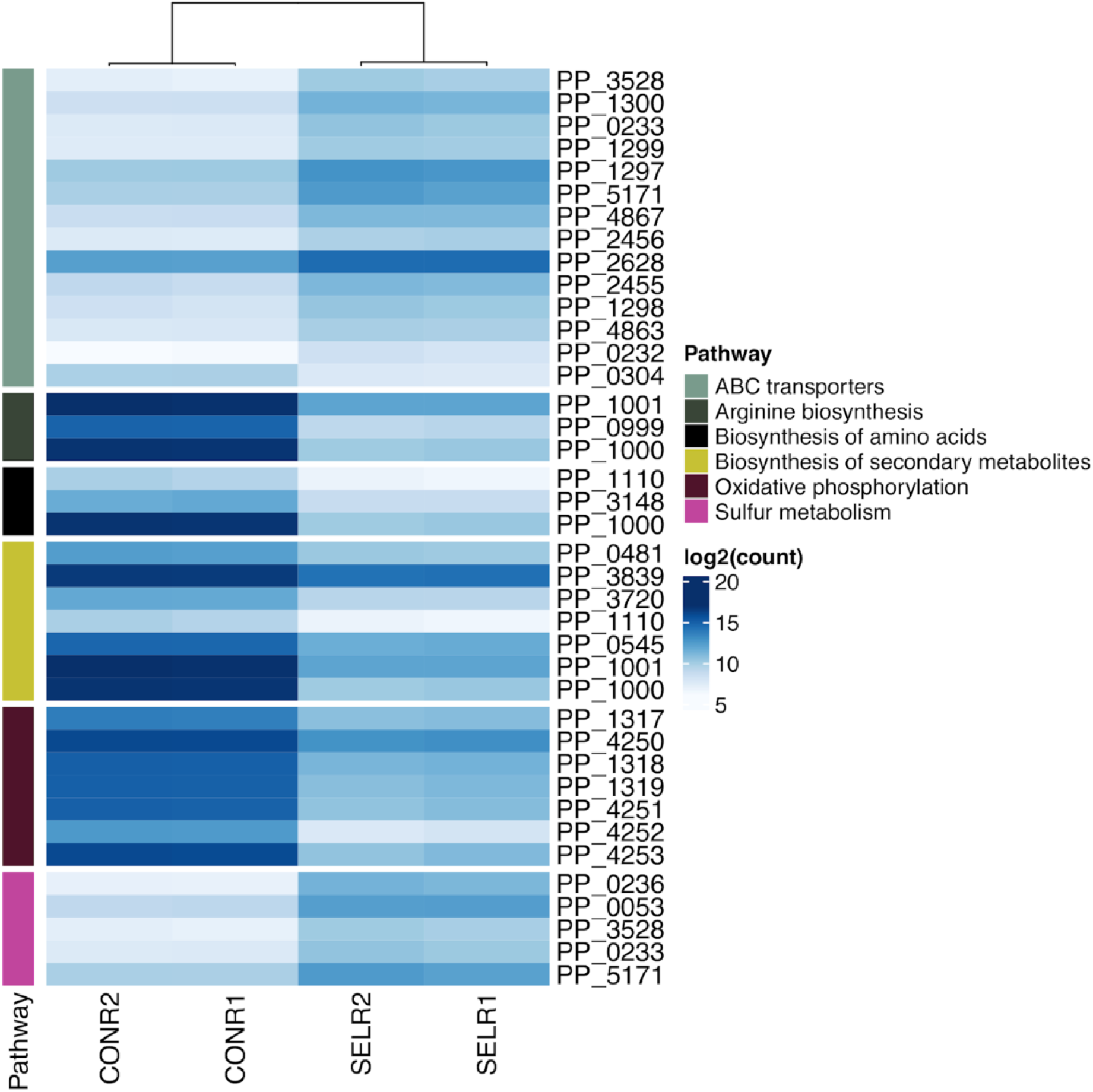
Heatmap representing selected pathways differentially expressed in the presence of selenite.

When investigating specific genes, the expression profile is consistent with a response to mitigate oxidative stress (see Supplementary file DEA.logFC2.csv). The most highly expressed genes include PP_3427, PP_3425 and PP_3426 encoding the multidrug efflux RND outer membrane protein OprN and the multidrug efflux RND transporter MexEF, which have been reported to be overexpressed in *P. putida* during the detoxification of formaldehyde (Roca *et al.*, 2008). Other highly expressed genes in presence of selenite encode the peroxidase LsfA, the alkylhydroperoxide reductase AhpCF (PP_2439 and PP_2440) and the semialdehyde dehydrogenase SadI (PP_3151). Some of the overexpressed genes involved in sulfur metabolism are the NAD(P)H-dependent FMN alkanesulfonate monooxigenase SsuEF (PP_0235 and PP_0236), the putative dimethyl sulfone monooxygenase SfnCE (PP_2772 and PP_2773) and the disulfide bond forming complex Dsb (PP_4235, PP_4236, PP_4237 and PP_4238). Out of the genes identified in the mutagenesis screening, only *sqr* and *pdo2* (PP_0052 and PP_0053), together with the last gene in the operon they form (PP_0054) were significantly overexpressed in response to selenite (Fig. 3). These genes are responsible for the fast phenotype when disrupted, which delay selenite reduction in the wild-type strain as their mutation leads to increase the rate of selenium production.

## Discussion

### Genes related to 2-ketoglutarate and glutamate metabolism participate in selenite reduction

*P. putida* possesses a robust central metabolism capable of reconfiguring the metabolic fluxes towards NADPH-producing reactions such as those catalyzed by the enzymes Zwf and GntZ (Nikel *et al.*, 2021). This cofactor is necessary for the action of the enzyme Glutathione reductase (Gor, PP_3819) and to maintain glutathione in its reduced form and thus fight against oxidative stress. Since glutathione is a key molecule, it is presumable that any changes in its biosynthesis may alter the cellular capacity to resist oxidative stress and carry out reduction processes. Mutagenesis with Tn5 transposons combined with a screening in the presence of selenite allowed to obtain two mutant strains (*sucA* and *D2HGDH*) deficient in the reduction of selenite which, according to its metabolic functions, affect the 2-ketoglutarate pool, a master regulator metabolite (Huergo and Dixon, 2015) and the starting point for glutamate and then glutathione biosynthesis. The first mutant has inactivated the *sucA* gene encoding the E1 subunit of 2-ketoglutarate dehydrogenase. This enzyme is responsible for transforming 2-ketoglutarate to succinyl-CoA in the Krebs cycle, so it could be presumed that *sucA* is an essential gene. However, as seen in Supp. Fig. S1, this mutant can grow very similarly to the wild-type strain, what could be explained in different ways. First of all, our experiments were performed in LB rich medium which can provide various carbon sources and synthesize essential metabolites from different metabolic steps. For example, Zhang *et al.* (2018) demonstrated that *P. putida* KT2440 could grow with L-lysine as a carbon source using glutamate as an intermediate and a combination of enzymes that can transform 2-ketoglutarate directly to succinate, that is, not via *sucA*. Second, in the absence of 2-ketoglutarate dehydrogenase, an alternative pathway to the Krebs cycle has been reported in several bacterial species (Green *et al.*, 2000, Xiong *et al.* 2014), so we hypothesized that in *P. putida* a similar cycle with glutamate, gamma-aminobutyric acid (GABA) and succinate semialdehyde could also be operating (Fig. 6).

**Figure 6.**
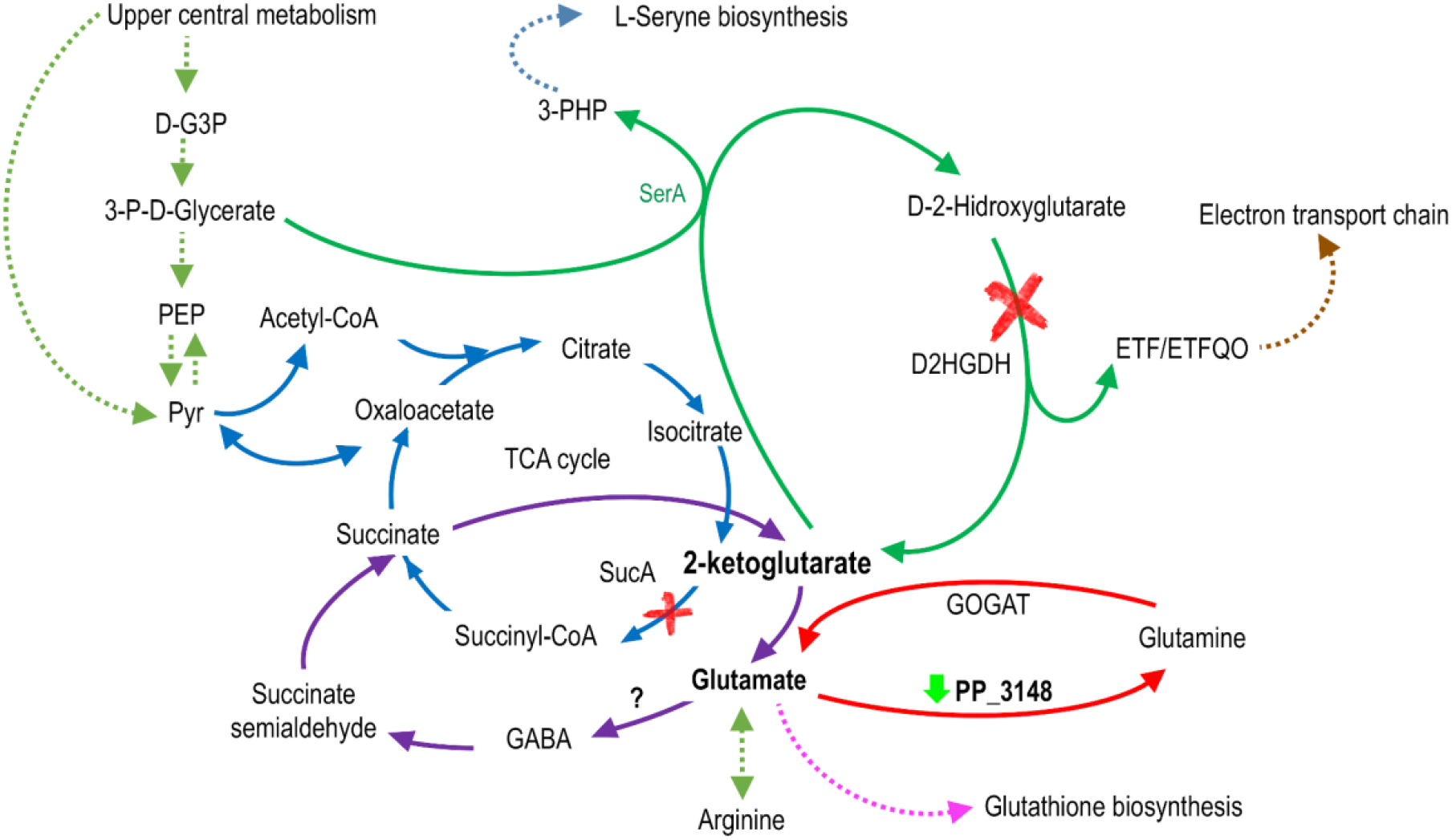
Reduced representation of 2-ketoglutarate and glutamate metabolism in *P. putida* **KT2440**. The mutation in the genes that encode the enzymes SucA and D2HGDH is represented with a red “X”. Mutation in these genes showed a deficient phenotype in selenite reduction. Both mutants correspond to enzymes related to the metabolism of 2-ketoglutarate, a key metabolite that is part of the Krebs cycle (blue) and is the starting point for the synthesis of glutamate and glutathione. Furthermore, recent studies connect this metabolite with serine synthesis (green). An alternative route for the synthesis of succinate (purple) that is not through the enzyme SucA is also represented in the figure. In the transcriptomic analysis, genes that were differentially expressed in the presence of selenite were observed. One of them is the PP_3148 gene that was overexpressed (downward green arrow) and encodes a glutamine synthetase.

According to the KEGG Pathway Database in the genome of *P. putida* KT2440, the enzymes responsible for transforming GABA to succinate semialdehyde (PP_0214 and PP_2799) and succinate semialdehyde succinate (PP_0213, PP_2488, PP_3151 and PP_4422) are present. However, an enzyme with glutamate decarboxylase activity responsible for transforming glutamate to GABA has not been described. Although there is no annotated enzyme for this reaction, in the literature, it has been described that several species of *Pseudomonas* are capable of synthesizing GABA (Dagorn *et al.*, 2013). In the context of this work, we consider that in a *sucA* mutant, an alternative pathway could be present (represented in purple in Figure 6), causing a reduction in glutamate availability which in turn could affect glutathione synthesis. In the wild-type strain glutamate synthesis is carried out from 2-ketoglutarate of the tricarboxylic acid cycle, serine synthesis or proline degradation (Gryder and Adams, 1969; Revelles *et al.*, 2005; Zhang *et al.*, 2016) and the pool of glutamate is used for biosynthesis of glutamine, glutathione and porphyrins such as cytochrome c and syroheme. However, in the *sucA* mutant according to the model shown in Fig. 6, 2-ketoglutarate and glutamate metabolism would require a reconfiguration of the metabolic network. Interestingly, in the transcriptomics experiments the genes involved in the transformation from 2-ketoglutarate to Succinyl-Co-A (i.e. *sucA*, *sucB* and *lpdA*), are underexpressed in the presence of selenite (Fig. S9), which supports the results obtained in mutagenesis. Other central metabolic genes such as succinate dehydrogenase and pyruvate carboxylase (Supp. Figure S9) are also underexpressed which supports the idea of the Krebs cycle and specifically 2-oxoglutarate is involved in the reduction of selenite. Possibly the presence of selenite generates oxidative stress affecting the pathways involved in glutathione synthesis, specifically in 2-oxoglutarate and glutamate metabolism. Underexpression of gene PP_3148 in transcriptomic analysis (Figs. 5 and 6), also supports this statement. PP_3148 encodes a glutamine synthetase responsible for the transformation of glutamate to glutamine, which again reaffirms the participation of 2-oxoglutarate and glutamate and their metabolism in selenite reduction.

A mutant in the D-2-hydroxyglutarate dehydrogenase (D2HGDH) also shows a deficient phenotype in the selenite reduction process (Fig. 6). We hypothesize that this mutation also affects the pool of 2-ketoglutarate and glutamate. D2HGDH catalyzes the reaction that converts D-2-hydroxyglutarate (D-2-HG) to 2-ketoglutarate. D-2-HG is a key player in the central metabolism of *Pseudomonas* including *P. putida* KT2440 (Zhang *et al.*, 2016). In these bacteria, the synthesis of L-serine is initiated by the enzyme SerA, which couples the dehydrogenation of D-3-phosphoglycerate to 3-phosphohydroxypyruvate and the reduction 2-ketoglutarate to D-2-HG (Zhang *et al.*, 2016) (see Fig. 6). Subsequently, 2-ketoglutarate is replenished from D-2-HG thanks to the D2HGDH activity (Zhang *et al.*, 2016). We hypothesize that by inactivating the enzyme D2HGDH, 2-ketoglutarate cannot be replaced from D-2-HG affecting the pool of 2-ketoglutarate and glutamate and as a consequence the biosynthesis of glutathione is also affected. This mutant not only allows us to strengthen our conclusions about the importance of 2-ketoglutarate and glutamate metabolism in selenite reduction but also allows us to make a connection with the synthesis of other amino acids, such as serine. (Fig. 6). The transcriptomic analysis also offers us evidence of variations in the expression of genes related to the metabolism of serine. The PP_1110 gene, underexpressed in the presence of selenite, encodes a serine acetyltransferase, an enzyme that links amino acid and sulfur metabolism.

The differentiated expression of genes related to the biosynthesis of arginine obtained in the presence of selenite in the RNA sequencing experiments (Figs. 4, 5 and Supp. Figure S5) also point to the participation of the metabolism of 2-oxoglutarate and glutamate in the reduction of selenite since the biosynthesis of arginine is carried out from glutamate or glutamine. As seen in Fig. 5 and Supp. Figure S5 in addition to glutamine synthetase (PP_3148; EC: 6.3.1.2), a carbamate kinase (PP_0999; EC: 2.7.2.2), an ornithine carbamoyltransferase (PP_1000; EC: 2.1.3.3), and an arginine deiminase (PP_1001; EC:3.5.3.6) involved in arginine biosynthesis are underexpressed in the presence of selenite.

### The response to oxidative stress is activated in the presence of selenite

The mutagenesis experiments allowed obtaining a deficient strain in the reduction of selenite that had the gene that encodes a glutathionyl-hydroquinone reductase (Gqr) truncated. This class of enzymes are widely distributed in bacteria, halobacteria, fungi, and plants and catalyze the reduction of glutathionyl-hydroquinones to hydroquinones (Green *et al.*, 2012). This enzyme belongs to the superfamily of Glutathione transferases that are involved in cellular detoxification against harmful xenobiotics and endobiotics (Allocati *et al.*, 2009). This reaction contributes to the regeneration of GSH (reduced glutathione) from GSSG (oxidized glutathione)((Fig. 7) which is undoubtedly involved in the oxidative stress response. In the Gqr mutant this replenishment is blocked, possibly directly impacting the GSH pool and its oxidative stress response. According to the functions described in the literature, there is no doubt that Gqr is essential for the cellular response to oxidative stress generated by selenite in *P. putida*.

**Figure 7.**
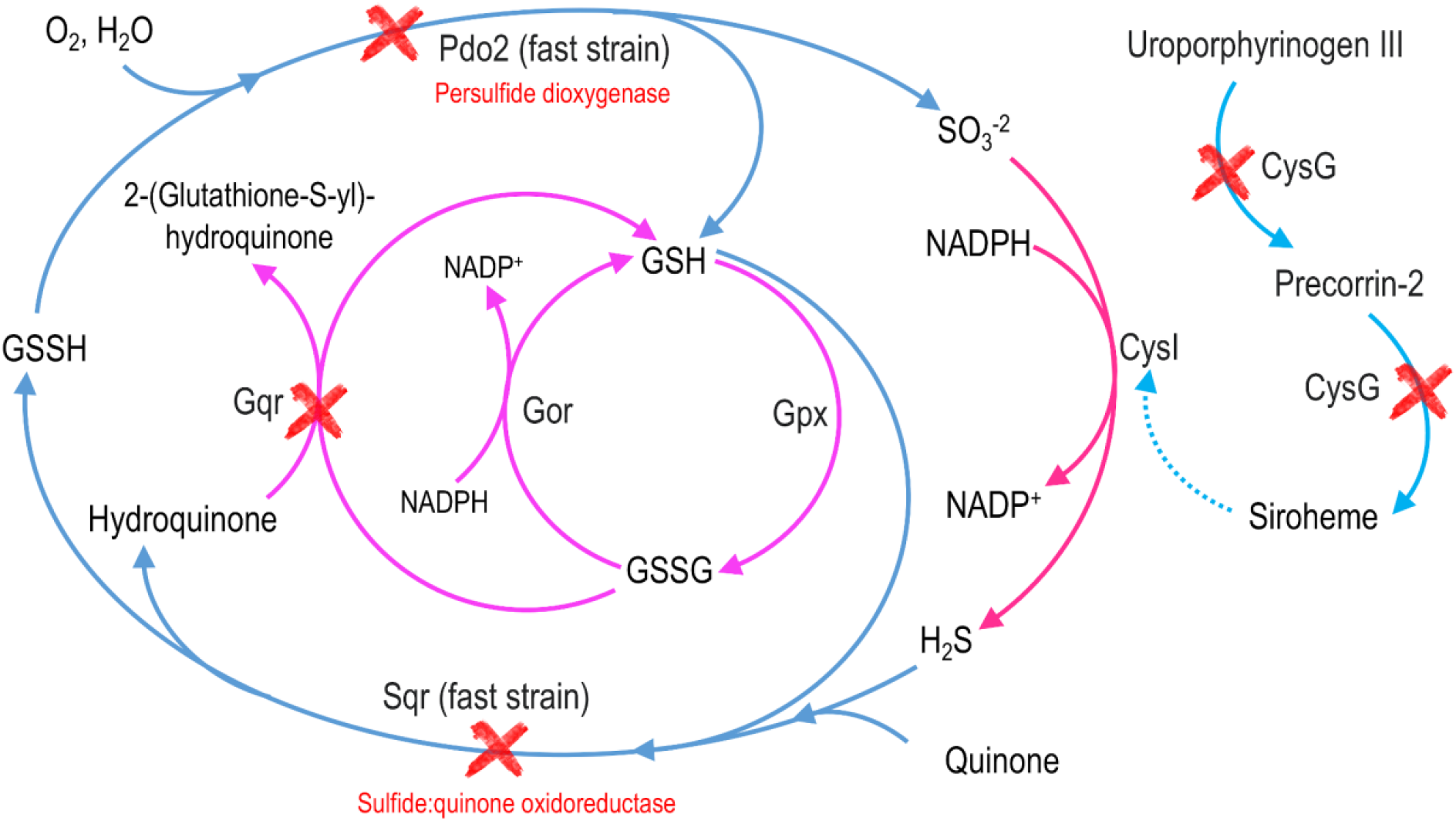
Reduced representation of sulfite and glutathione metabolism in *P. putida* **KT2440**. The genes that were truncated in the mutagenesis experiments are marked with a red “X”. Mutation in genes *cysG* and *gqr* showed a deficient phenotype, while mutation in genes *sqr* and *pdo2* showed a fast phenotype in selenite reduction. Fast strains are truncated in genes that recycle H_2_S (and maybe H_2_Se) to SO_3_^−2^ (and maybe SeO_3_^−2^), suggesting that sulfide:quinone oxidoreductase and persulfide dioxygenase constitute a pathway that competes with the reduction of selenite to elemental selenium. On the other hand, the mutation in *cysG* suggests an impairment in the activity of sulfite reductase (CysI). A mutant in the PP_2370 gene (which forms an operon with *cysI*) was also obtained during mutagenesis, reinforcing the idea that sulfite reductase activity is important for selenite reduction. The mutation in *gqr* also indicates the importance of glutathione metabolism in selenite reduction.

The transcriptomic analysis also reflected the overexpression of other genes related to the response to oxidative stress namely the peroxidase LsfA, the alkylhydroperoxide reductase AhpCF (PP_2439 and PP_2440) and the semialdehyde dehydrogenase SadI (PP_3151)(Fig. 3). Kaihami *et al.* (2014) reported that LsfA from the opportunistic pathogen *Pseudomonas aeruginosa* is endowed with thiol-dependent peroxidase activity that protects the bacteria from H_2_O_2_. In *Pseudomonas fluorescens* it was also shown that this gene encodes an enzyme that has peroxidase and thioredoxin activity (Liu *et al.*, 2015). Moreover, the alkylhydroperoxide reductase AhpCF proved to be critical for survival of biofilm bacteria in *P. aeruginosa* in the presence of hydrogen peroxide (Panmanee and Hassett, 2009). The importance of AhpCF in resistance to oxidative stress has been well documented in several bacterial genera (Rocha and Smith, 1999; Wan *et al.*, 2019; Feng *et al.* 2020). Therefore, the overexpression of the genes encoding LsfA and AhpCF in the presence of selenite, confirms that this oxyanion induces the response to oxidative stress in *P. putida*.

### Sulfur metabolism is involved in selenite reduction

The nonspecific activity of some oxidoreductases on selenite is a controversial issue. It has been suggested that due to the structural similarity and chemical reactivity of species such as SO_3_^−2^ and SeO_3_^−2^, some enzymes can act on them in a non-specific way. In the case of selenite, due to the proximity in the periodic table of sulfur and selenium, it is presumable that enzymes such as sulfite reductase (responsible for transforming sulfite to hydrogen sulfide; see Fig. 7) can recognize selenite. In species of *Alcaligenes faecalis*, it was reported that selenite reduction occurs through the activity of proteins such as sulfite reductase and thioredoxin reductase (Wang *et al.*, 2018). Our results indicate that sulfur metabolism is important for selenite reduction in *P. putida* KT2440. We isolated four mutants whose disrupted genes are related to sulfite reductase activity (mutants ID RF30, RF31, JJ3, and JJ7; Table 1). These mutants contain insertions in PP_2370 (unknown protein forming an operon with *cysI* encoding the Heme protein sulfite reductase, beta subunit) and *cysG* genes (PP_3999, encodes a Uroporphyrin-III C-methyltransferase) (Table 1). It is important to highlight that in the screening we isolated three mutants affected in *cysG* (Table 1, mutants ID RF30, RF31 and JJ3), which suggests that this gene is crucial for selenite reduction. It has previously been reported in *E. coli* that the sulfite reductase requires syroheme as a cofactor (Wu *et al.*, 1991). Therefore, we hypothesize that the activity of the uroporphyrin-III C-methyltransferase is required for the synthesis of siroheme, a cofactor required for the action of the product encoded by *cysI*. Then, the hydrogen selenide produced by the activity of CysI may react with ambient oxygen in an abiotic manner, generating elemental selenium and superoxide as described in (Kessi and Hanselmann, 2004). We propose that the *cysI-cysG* system performs at least partially the reduction of selenite to hydrogen selenide, following the pathway represented in Fig. 7 for sulfur metabolism.

The involvement of sulfur metabolism in selenite reduction is supported by two other mutants obtained in this study: *sqr* (PP_0053), which encodes a sulfide-quinone oxidoreductase and *pdo2*, which encodes a persulfide dioxygenase and whose function already has been experimentally demonstrated (Shibata and Kobayashi, 2006; Sattler *et al.*, 2015). It is worth noting that we also isolated a mutant in the gene encoding for the putative transcriptional activator of the operon containing these genes (PP_0051). As shown in figure 7, the Sqr/Pdo2 system is responsible for recycling part of the H_2_S (and maybe H_2_Se) to SO_3_^−2^ (and maybe SeO_3_^−2^). The Sqr/Pdo2 system, which is opposed to the activity of the sulfite reductase, has been proposed as a mechanism responsible for maintaining the concentration of H_2_S (and maybe H_2_Se) in equilibrium between its formation and oxidation (Sattler *et al.*, 2015). In the first step (catalyzed by Sqr, Fig. 7), the use of H_2_S is coupled with quinone and GSH to produce GSSH and hydroquinone. The sulfide-quinone oxidoreductase activity of Sqr has been described in *P. putida* KT2440 (Shibata and Kobayashi, 2006) and its deletion led to a decrease in the intracellular catalase and ubiquinone-H2 oxidase activity. In the second step, the enzyme Pdo2 oxidises glutathione persulfide (GSSH) to sulfite and GSH (Sattler *et al.*, 2015). The overexpression of these genes in the presence of selenite suggests that processes which may be considered antagonistic – selenite reduction and its prevention by, respectively, deficient and fast genes – act at the same time, possibly to ensure the appropriate balance of activities and usage of cofactors. As shown in Figure 2 and Supp. Video S1, the *sqr* and *pdo2* mutants metabolize selenite more rapidly than the wild-type strain, which suggests that blocking these two metabolic steps increases the availability of H_2_Se for its subsequent reduction to Se°. During this cyclical process (SeO_3_^−2^→H_2_S→SeO_3_^−2^) mediated by Sqr/Pdo2, there is no net consumption of GSH. The identification of the enzymes Sqr and Pdo2 as targets to increase the production rate of selenium nanoparticles is of interest for the development of biotechnological tools.

The participation of sulfur metabolism in selenite reduction was also evidenced in transcriptomics experiments. Several genes related to sulfur metabolism were overexpressed including the NAD(P)H-dependent FMN alkane sulfonate 17onooxygenase SsuEF, the putative dimethyl sulfone monooxygenase SfnCE and the disulfide bond forming complex Dsb (Fig. 5 and Supp. Figure S4). Genes related to disulfide bond formation are very consistent with a previous proposal for the biological reduction of selenite *in E. coli* (Kessi and Hanselmann, 2004). In this model, a sequence of transformations with the formation of molecules with S-Se bonds is proposed. Transcriptomic experiments also revealed that exposure to selenite triggered the expression of membrane associated proteins associated to sulfur (Fig. 5). It was possible to observe in the presence of selenite the overexpression of transporters related to the uptake of sulfate (PP_5171) and the sulfur compound taurine (PP_0232, PP_0233) (Fig. 5 and Supp. Figure S4). Previous studies provided biochemical information that both selenite and selenate could be transported in *E. coli* K-12 through a sulfate transporter (Lindblow-Kull *et al.*, 1985). Lusa *et al.* (2017) through a proteomic approach studied the possible selenite transport systems in two environmental isolates of *Pseudomonas*. The authors suggest two different transport mechanisms for Se(IV) uptake in these *Pseudomonas* sp. strains; a low affinity transport system up-regulated by NO_3_^−^/NO_2_^−^/SO_4_^2−^ and a distinct SeO_3_^2−^ regulated transport system. Considering this information, it is presumable that the transporter PP_5171 may be involved in selenite uptake in *P. putida* KT2440, reaffirming the importance of sulfur metabolism including transport systems.

### Other genes involved in selenite reduction

In the mutagenesis experiments, a deficient strain in selenite reduction was obtained with an insertion in the gene *ccmF*, which encodes a holocytochrome C synthetase and is part of the well-known cytochrome c maturation system. All the components of the Ccm system are located in the cytoplasmic membrane, which is responsible for the covalent attachment of haem to the CXXCH motif of apocytochrome in the periplasm (Cianciotto *et al.*, 2005). Mutations in genes of the Ccm system have shown a great variety of phenotypes in bacteria, some of them with opposite effects, which means that its function is still unclear. For example, in *Pseudomonas fluorescens* (Gabala *et al.*, 1996; Gaballa *et al.*, 1998), *Pseudomonas aeruginosa* (de Chial *et al.*, 2006) and other bacterial genera (Pearce *et al.*, 1998), mutations in *ccm* genes resulted in reduced production and uptake of siderophores. However, in *P. putida* GB-1, a reduction in pyoverdine production was not observed in different *ccmF* mutants (de Vrind *et al.*, 1998). This study demonstrated that in *P. putida* GB-1 mutants in *ccmF* did not have cytochrome oxidase activity, did not contain c-type cytochromes, and was deficient in the oxidation of Mn^2+^. Cianciotto *et al.* (2005) proposed that low levels of cytosolic haem are produced in *ccm* mutants, which prevents the maturation of several enzymes, including those involved in the biosynthesis of siderophores or in the resistance of oxidative stress. This relationship between the deficiency of the *ccm* genes and the impairment of the response to oxidative stress could explain the phenotype observed in the mutant *ccmF* concerning the reduction of selenite. Transcriptomic analysis also revealed that the presence of selenite led to the undeexpression of two operons related to cytochromes activity and oxidative phosphorylation (Fig. 5 and Supp. Figure S10). The PP_1317-PP_1319 operon encodes a ubiquinol-cytochrome c reductase and the PP_4250-PP_4253 operon encodes a cbb3-type cytochrome c oxidase. It is well known that oxidative phosphorylation has the function of oxidizing nutrients in order to produce adenosine triphosphate (ATP). That these operons are underexpressed in the presence of selenite could reflect the metabolic failures and damage to cell function that the presence of this oxyanion can generate.

During mutagenesis, we isolated other strains with Tn5 disruptions in genes encoding proteins related to the cell membrane, namely *msbA* (a lipid transporter ABC ATPase), *wzy* (an O-antigen polymerase) and PP_4799 (a muranoyltetrapeptide carboxypeptidase). The *msbA* and *wzy* genes form an operon in *P. putida* KT2440 and by homology with studies carried out in *P. aeruginosa* (Ghanei *et al.*, 2007; Huszczynski *et al.*, 2020) and other bacteria (Singh *et al.*, 2016; Padayatti *et al.*, 2019; Wiseman *et al.*, 2021) it is possible that in *P. putida* both genes are related to lipopolysaccharides (LPS) trafficking and assembly. Specifically, MsbA is an essential ATP-binding cassette transporter in Gram-negative bacteria that flips lipid A with or without core sugars from the cytoplasmic leaflet to the periplasmic leaflet of the inner membrane (Padayatti *et al.*, 2019). In *P. aeruginosa*, it was shown that *msbA* is an essential gene and that the activity of MsbA can be stimulated by lipid A linked to complete core oligosaccharides (Ghanei *et al.*, 2007). The gene *wzy* encodes a polymerase responsible for the biosynthesis of polysaccharides, including LPS (Wiseman *et al.*, 2021). Finally, a mutant in the PP_4799 gene encoding a muramoyltetrapeptide carboxypeptidase (also known as LD-Carboxypeptidase) was obtained. Homologs in other bacterial genera indicate that these enzymes can cleave amide bonds that link an L-aminoacid to a C-terminal D-aminoacid, and a role for these enzymes in peptidoglycan recycling has been proposed (Korza and Bochtler, 2005). Based on the results obtained in this study it is difficult to assign a clear function to these genes (i.e., *msbA*, *wzy* and PP_4799) in the metabolism of selenium. It is likely that they participate in the transport of selenite/selenium in the cell or rather they are a response at the membrane level as a result from selenite reduction and presence of selenium nanoparticles.

Transcriptomic experiments also revealed that exposure to selenite triggered the expression of membrane associated proteins (Fig. 5). It was possible to observe in the presence of selenite the overexpression of transporters related to the uptake of amino acids (operon PP_1297, PP_1298, 1299 and PP_1300) and ribose (PP_2456, PP_2455). The differences in expression patterns in membrane proteins can be explained considering that selenite reduction leads to the accumulation of extracellular selenium nanoparticles, which could lead to a significant rearrangement of the membrane proteome.

## Conclusion

This work provides the first molecular insights of selenium metabolism in *P. putida* KT2440. This bacterium has a specialized central metabolism and enzymatic machinery for producing high reducing power. This characteristic makes *P. putida* an excellent bacterial model for studying chemical biotransformations that involve a reduction process, such as transforming selenite into selenium. Our results suggest that the reduction process is carried out by an interconnected pathway involving sulfur metabolism (represented by genes *cysG, sqr, pdo2, sqrR, ssuEF and sfnCE*), 2-oxoglutarate/glutamate metabolism (represented by genes *sucA, D2HGDH, PP_3148*) and oxidative stress response (represented by genes*,Gqr, lsfA, ahpCF* and *sadI*). Genes *sucA*, D2HGDH and PP_3148 encode central metabolic enzymes related to the 2-ketoglutarate metabolism, a key molecule and the starting point for the synthesis of glutamate and glutathione, so that these central metabolic reactions could also be related to the response to oxidative stress. Thus, considering that sulfur metabolism is also interconnected with glutathione metabolism (Fig. 7), we propose that in *P. putida* KT2440, the reduction of selenite is carried out through the metabolism of inorganic sulfur (SO_3_^−2^→H_2_S) and the response to oxidative stress mediated by glutathione (Fig. 8). This study not only sheds light on the metabolism of selenium in *P. putida* but also (i) reports the involvement of genes that had not been related to the metabolism of selenium in any other bacterial genus and (ii) and visualizes genes of biotechnological interest (*sqrR, pdo2* and *sqr*) whose suppression generates the production of selenium nanoparticles at a higher rate than the wild-type strain. The data provided in this study brings us closer to understanding the metabolism of selenium in bacteria, and to the development of biotechnological tools that can be used to recover this element from the environment.

**Figure 8.**
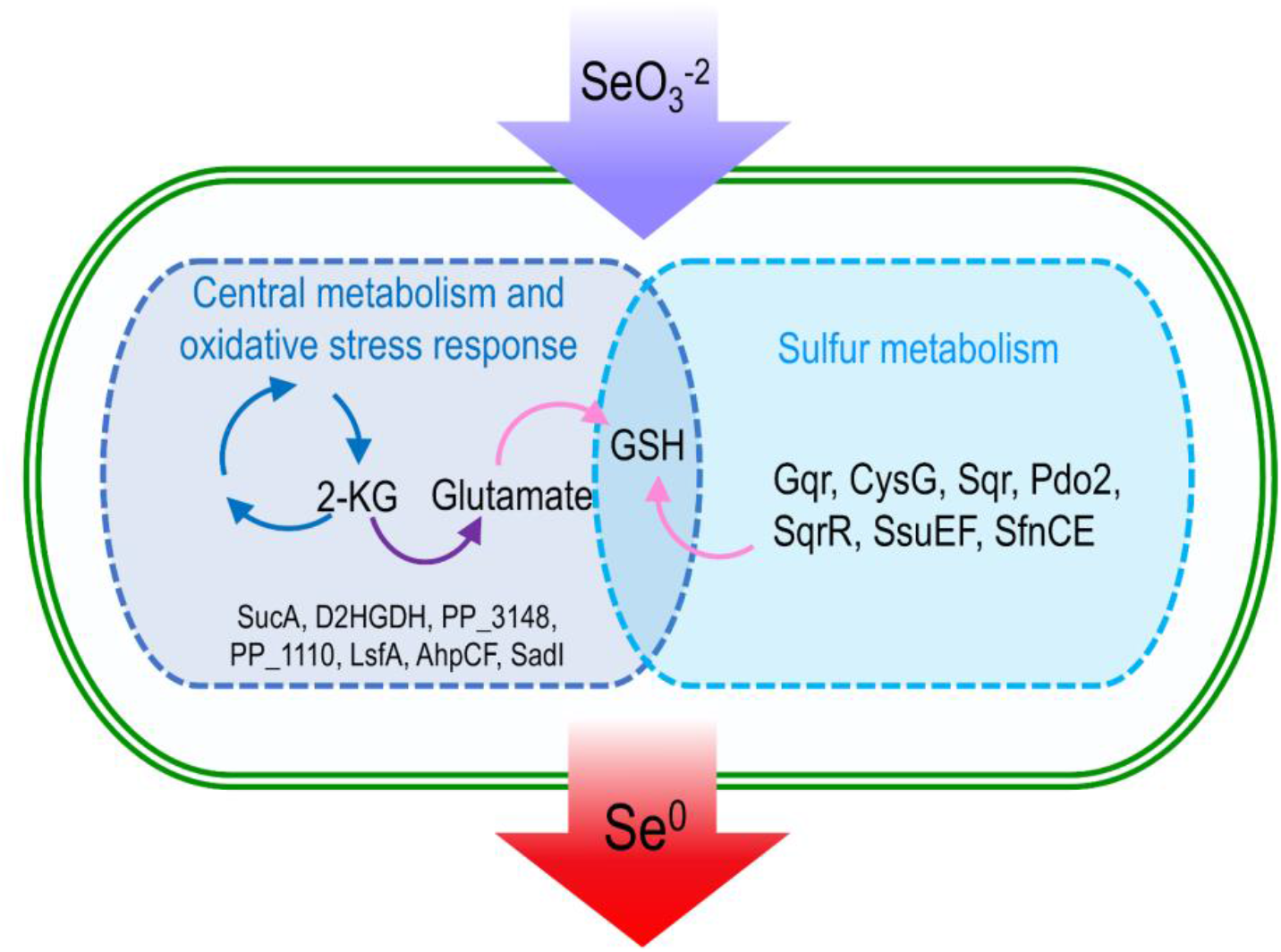
Molecular actors involved in selenite reduction in *P. putida* **KT2440**. Enzymes of central metabolism, oxidative stress response and sulfur metabolism participating in the reduction of selenite to selenium nanoparticles. Both steps converge in the production of glutathione.

## Methods

### Bacterial strains, plasmids, culture media and growth conditions

*P. putida* KT2440 was cultured aerobically in LB broth at 30°C with orbital shaking at 150 rpm. *E. coli* strains were cultured in LB broth at 37°C with orbital shaking at 200 rpm. *E. coli* CC118λpir (pBAM1) (Martínez-García *et al.*, 2011) medium was supplemented with ampicillin (100 μg/mL) and kanamycin (50 μg/mL) while *E. coli* HB101 (pRK600) medium was supplemented with chloramphenicol (30 μg/ml). All chemicals were purchased from Sigma-Aldrich (UK).

### Mutagenesis with pBAM1

Mutagenesis of *P. putida* KT2440 by random insertion of miniTn5 transposons was conducted as described in (Martínez-García *et al.*, 2011). Briefly, pBAM was transferred to *P. putida* via triparental mating using *E. coli* CC118λpir (pBAM1) as a donor and *E. coli* HB101 (pRK600) as a helper. Each strain was grown independently overnight in LB broth and the cell pellet of 1 mL of each culture was washed with 1 mL of LB three times before resuspending in 100 μL of LB and mixed with the other strains. 100 μL of the mixture were deposited on a 0.22 μm nitrocellulose filter on a LB plate and incubated overnight at 30°C. After conjugation the cells on the filter were recovered by resuspending in 1 mL of LB and plated in selective media containing *Pseudomonas* Isolation Agar (Gifco; UK), kanamycin and 1 mM sodium selenite. Plates were incubated at 30°C until colonies achieved an observable phenotype. The location of the Tn5 insertions was determined using arbitrary PCR of genomic DNA obtained from cell colonies as described previously (Das *et al.*, 2005; Martínez-García *et al.*, 2011). PCR products were subject to Sanger sequencing and the resulting sequences were subject of blast (Altschul *et al.*, 1990) against the *P. putida* KT2440.

### Cloning of genes from *P. putida* KT2440 for complementation tests

For mutant complementations the protocol described by Montenegro *et al.* (2021) was used. Briefly the DNA sequence encoding the respective gene for each mutant (see Table 1) of P. *putida* KT2440 was amplified directly from the chromosome of the bacterium with their corresponding primers (see Supplementary Table S3). The resulting amplicon was digested (see supplementary Table S3), and further ligated to the polylinker of the expression plasmid pSEVA438 as an either EcoRI/HindIII or BamHI/HindIII fragment (See Supplementary Table S3) and subsequently transformed and propagated in *E. coli* DH5α. To verify the ligation, we replated colonies in fresh media with streptomycin (Sm 100 μg ml^−1^); a colony PCR was performed with the respective primers for each gene. After verifying the ligation of the plasmid we transformed the construct into the deficient strains (See Table 1). Deficient strains were also transformed with the empty backbone (pSEVA438) as a control. To establish if the genetic complementation of the mutant was enough for acquiring the wild-type phenotype again, we grew preinocula of the mutant strains complemented and with the empty backbone (pSEVA438) overnight in Luria-Bertani (LB) as described above. Optical density (OD) was measured and adjusted to 0.05 in fresh media supplemented with selenite (1 mM), Sm (100 μg ml^−1^) and 3-methylbenzoate (3MB, 0.5 mM). Strains were incubated with orbital shaking at 175 rpm at 30 °C. Phenotype was analyzed after 24 h of incubation.

### Growth curves of the mutants during selenite reduction

The effect of the mutations on *P. putida* KT2440 cultured in the presence or absence of selenite was assessed by monitoring growth kinetics at 600 nm (OD600) of the wild-type and mutant strains in 96 well microtiter plates (Surface Nunclon ™ Roskilde, Denmark) using a Sinergy H1 Hybrid Multi-Mode microplate reader (Biotek, USA). For this, cultures of each strain were diluted to an initial optical density at 600 nm (OD600) of approximately 0.05 in fresh medium with 0 and 1 mM sodium selenite in a volume of 200 μL per well. Microplates were incubated at 30°C for 24 hr with continuous orbital shaking and measurement of optical density every 10 minutes. Each growth curve consisted of 3 biological replicates. Photos of the plates were taken at 24 h of the experiment to record the coloration of the medium.

### Determination of residual selenite amount by FAAS (Flame Atomic Absorption Spectroscopy)

Selenite reduction was recorded by analysis of residual selenite in the culture medium. For this, 3 L Erlenmeyer flasks with 500 mL of LB supplemented with 1 mM selenite were inoculated with *P. putida* KT2440 (initial OD of 0.05). The mutant strains were grown at 30°C and 150 rpm for 24 h. For deficient mutants, 15 mL aliquots of the cell suspension were taken at 0 h and 24 h during bacterial growth, while for mutants with the capacity to produce selenium more rapidly, samples were taken between 0 h and 20 h with intervals of 2.5 h. Then, its OD600 was measured, and the cells were collected by centrifugation (10 min, 4000 × g, 4°C). The supernatant was then carefully transferred to a new 15 mL tube and stored at −25°C until further analysis. For analytical determinations, supernatants were filtered on mixed cellulose ester syringe filters (0.20 μm, ADVANTEC®, USA) and analyzed using an atomic absorption spectrometer (AA240FS + 240Z, Varian Inc., USA). The wavelength (nm) used for selenium was 196,026. Selenite solutions were prepared from a 1,000 mg/L standard solution (Tritrisol-Merck, Darmstadt, Germany). Appropriate dilutions were made to prepare the standards, which were stored in polyethylene flasks under refrigeration. The average of three replicates is presented in the bars and the error was calculated as the standard deviation of the results.

### RNA sequencing experiments

*P. putida* KT2440 was precultured overnight in LB medium, and the culture was then diluted 100-fold in the same medium and grown to OD600 of 0.2. Samples were then either cultured further without additional modifications or treated with 1 mM selenite for 1 hour. Transcription in each culture was halted by spiking a high concentration of rifampicin (200 μg/mL). Total RNA was isolated from 1 ml of cells from each culture using a miRNeasy kit (Qiagen, Germany) with some modifications. The collected pellets were resuspended into 0.1 mL Tris–HCl (pH 7.5) containing lysozyme (2 mg/mL) and incubated for 10 min at 37°C. The lysate was processed according to manufacturer’s instructions. RNase-free DNase (Qiagen, Germany) treatment was performed during the RNA isolation procedure to eliminate residual DNA and the quality of isolated RNAs was evaluated using a 2100 Bioanalyzer System (Agilent, USA). To obtain transcriptional profile at the genome-wide level, libraries of cDNA with an average size of 200 nucleotides obtained from two independent replicates per condition were sequenced using a HiSeq 2000 platform (Illumina, USA) by Novogene Corporation (China). The raw data was uploaded to GEO with accession number GSE214391.

The DESeq2 package version 1.34.0 was used for normalisation and differential expression analysis between control and treatment samples following mapping and counting, with effect size shrinkage using the ‘apeglm’ approach (Zhu *et al.*, 2019) (processed normalised counts are compiled in the Supplementary File ‘Selenite.RawCounts.csv’). Gene annotation was carried out using the tool of the DESeq2 package and the appropriate file (gtf) from the assembly (GCA_000014625.1). Genes with a log-Fold Change > 2 and adjusted p-value <0.05 were considered differentially expressed (DEGs). DEGs were functionally classified using the goseq R package version 1.46 (Young *et al.*, 2014) and the Kyoto Encyclopedia of Genes and Genomes (KEGG) database (http://www.genome.jp/kegg/) (Kanehisa and Goto, 2000). Go annotation and KEGG classifications were downloaded from the *Pseudomonas* Community Annotation Project (PseudoCAP).

## Supporting information

Supplementary file DEA.logFC2.csv

## Author Contributions

JJ and MC conceived and designed the experiments. RA, SV, RM, PF, DR-G, RF, MS and JK performed the experiments. RA, SM, JJ, MS, JK and MC analyzed the data. PF, JIJ and MC contributed reagents or materials, or analysis tools. RA, SM, MC and JJ wrote the paper. All authors reviewed and approved the final version of the manuscript.

## Acknowledgements

This work was supported by the Vicerrectory of Research of the Universidad of Costa Rica (809-B5-A68). The authors also acknowledge to CENIBiot and CELEQ for financial support. JJ, acknowledges the support received from the Biology and Biotechnology research council (Grant no. BB/T011289/1 from the ERA-Cobiotech Programme of the EU). JJ and MS acknowledge the support from the European Community’s H2020 Programme (H2020-FNR-11-2020: SECRETED, Grant No. 101000794). SMM is the recipient of a President’s Scholarship from Imperial College London.

