## Supplementary information and figures for "Production of selenium nanoparticles occurs through an interconnected pathway of sulfur metabolism and oxidative stress response in *Pseudomonas putida* KT2440"

#### Supplementary information for

### Selenite reduction in *Pseudomonas putida* KT2440 occurs through an interconnected pathway involving inorganic sulfur metabolism and oxidative stress response

Roberto Avendaño[1], Said Muñoz-Montero[2], Diego Rojas-Gätjens[1], Paola Fuentes[3,4], Sofía Vieto[1], Rafael Montenegro[1], Manuel Salvador[5], Rufus Frew[6], Juhyun Kim[7], Max Chavarría[1,3,8*] and Jose I. Jiménez[2*]

[1]Centro Nacional de Innovaciones Biotecnológicas (CENIBiot), CeNAT-CONARE, 1174-1200 San José (Costa Rica). [2]Department of Life Sciences, Imperial College London, South Kensington campus, London SW7 2AZ (United Kingdom). [3]Escuela de Química, Universidad de Costa Rica, 11501-2060 San José (Costa Rica). [4]Centro de Electroquímica y Energía Química (CELEQ), Universidad de Costa Rica, 11501-2060 San José (Costa Rica). Biotechnology Applications, IDENER Research & Development, Earle Ovington 24 Nave 8-9, Seville 41300 (Spain). [5]Department of Chemistry, University of Leicester, Leicester LE1 7RH, (United Kingdom). [6]School of Life Sciences, BK21 FOUR KNU Creative BioResearch Group, KNU Institute for Microorganisms, Kyungpook National University, Daegu 41566 (Republic of Korea). [7]Centro de Investigaciones en Productos Naturales (CIPRONA), Universidad de Costa Rica, 11501-2060 San José (Costa Rica).

---

*Correspondence to: Jose Jimenez
Department of Life Sciences
Imperial College London
South Kensington Campus
London, SW7 2AZ
United Kingdom
Phone: (+44) 207594 3895

ORCID ID https://orcid.org/0000-0002-2700-9473

*Correspondence to: Max Chavarría
Escuela de Química & Centro de Investigaciones en Productos Naturales (CIPRONA)
Universidad de Costa Rica
Sede Central, San Pedro de Montes de Oca
San José, 11501-2060, Costa Rica
Phone (+506) 2511 8520. Fax (+506) 2253 5020

ORCID ID https://orcid.org/0000-0001-5901-3576

### Supplementary Tables

**Supplementary Table S1**. List of overexpressed Gene Ontology functions

| category | pvalue | numDEInCat | numInCat | term | ontology |
|---|---|---|---|---|---|
| GO:0097428 | 5.65E-06 | 3 | 3 | protein maturation by iron-sulfur cluster transfer | BP |
| GO:0016226 | 0.00010214 | 3 | 6 | iron-sulfur cluster assembly | BP |
| GO:0042908 | 0.00103939 | 2 | 4 | xenobiotic transport | BP |
| GO:0045454 | 0.00398757 | 3 | 19 | cell redox homeostasis | BP |
| GO:0042952 | 0.01228073 | 2 | 10 | beta-ketoadipate pathway | BP |
| GO:0000302 | 0.01607751 | 1 | 1 | response to reactive oxygen species | BP |
| GO:0044571 | 0.01669936 | 1 | 1 | [2Fe-2S] cluster assembly | BP |
| GO:0015734 | 0.0171253 | 1 | 1 | taurine transport | BP |
| GO:0046306 | 0.01776922 | 1 | 1 | alkanesulfonate catabolic process | BP |
| GO:0051259 | 0.01789011 | 1 | 1 | protein complex oligomerization | BP |
| GO:0006572 | 0.03283525 | 1 | 2 | tyrosine catabolic process | BP |
| GO:0008652 | 0.03415367 | 1 | 2 | cellular amino acid biosynthetic process | BP |
| GO:0009072 | 0.03435046 | 1 | 2 | aromatic amino acid family metabolic process | BP |
| GO:0019303 | 0.03502243 | 1 | 2 | D-ribose catabolic process | BP |
| GO:0006865 | 0.03688803 | 3 | 43 | amino acid transport | BP |
| GO:0019477 | 0.04917817 | 1 | 3 | L-lysine catabolic process | BP |
| GO:0006559 | 0.04971006 | 1 | 3 | L-phenylalanine catabolic process | BP |
| GO:0016020 | 0.00079977 | 5 | 44 | membrane | CC |
| GO:0042597 | 0.0187574 | 4 | 61 | periplasmic space | CC |
| GO:0015185 | 0.00026477 | 2 | 2 | gamma-aminobutyric acid transmembrane transporter activity | MF |
| GO:0015407 | 0.00081082 | 2 | 3 | ABC-type monosaccharide transporter activity | MF |
| GO:0047569 | 0.00179769 | 2 | 4 | 3-oxoadipate CoA-transferase activity | MF |
| GO:0015562 | 0.00603957 | 3 | 24 | efflux transmembrane transporter activity | MF |
| GO:0005198 | 0.0072653 | 2 | 8 | structural molecule activity | MF |
| GO:0051536 | 0.00780946 | 3 | 25 | iron-sulfur cluster binding | MF |
| GO:0051920 | 0.008249 | 2 | 8 | peroxiredoxin activity | MF |

| | | | | | |
|---|---|---|---|---|---|
| GO:0016887 | 0.00876814 | 6 | 108 | ATP hydrolysis activity | MF |
| GO:0004601 | 0.01095317 | 2 | 10 | peroxidase activity | MF |
| GO:0015424 | 0.01480005 | 2 | 11 | ABC-type amino acid transporter activity | MF |
| GO:0004485 | 0.01599694 | 1 | 1 | methylcrotonoyl-CoA carboxylase activity | MF |
| GO:0008785 | 0.01607751 | 1 | 1 | alkyl hydroperoxide reductase activity | MF |
| GO:0004334 | 0.01656069 | 1 | 1 | fumarylacetoacetase activity | MF |
| GO:0047075 | 0.01698453 | 1 | 1 | 2,5-dihydroxypyridine 5,6-dioxygenase activity | MF |
| GO:1901681 | 0.01702636 | 1 | 1 | sulfur compound binding | MF |
| GO:0050498 | 0.01711491 | 1 | 1 | oxidoreductase activity, acting on paired donors, with incorporation or reduction of molecular oxygen, with 2-oxoglutarate as one donor, and the other dehydrogenated | MF |
| GO:0004747 | 0.017234 | 1 | 1 | ribokinase activity | MF |
| GO:0004490 | 0.01739332 | 1 | 1 | methylglutaconyl-CoA hydratase activity | MF |
| GO:0047507 | 0.01741379 | 1 | 1 | (deoxy)nucleoside-phosphate kinase activity | MF |
| GO:0015411 | 0.01743935 | 1 | 1 | ABC-type taurine transporter transporter activity | MF |
| GO:0016034 | 0.01770356 | 1 | 1 | maleylacetoacetate isomerase activity | MF |
| GO:0008752 | 0.01776922 | 1 | 1 | FMN reductase activity | MF |
| GO:0102039 | 0.01781964 | 1 | 1 | alkylhydroperoxide reductase activity | MF |
| GO:0001671 | 0.01789011 | 1 | 1 | ATPase activator activity | MF |
| GO:0047134 | 0.03126207 | 1 | 2 | protein-disulfide reductase (NAD(P)) activity | MF |
| GO:0004777 | 0.03251367 | 1 | 2 | succinate-semialdehyde dehydrogenase (NAD+) activity | MF |
| GO:0003871 | 0.03410238 | 1 | 2 | 5-methyltetrahydropteroyltriglutamate-homocysteine S-methyltransferase activity | MF |
| GO:0008961 | 0.03446826 | 1 | 2 | phosphatidylglycerol-prolipoprotein diacylglyceryl transferase activity | MF |
| GO:0005506 | 0.03490484 | 3 | 42 | iron ion binding | MF |
| GO:0052873 | 0.0350432 | 1 | 2 | FMN reductase (NADPH) activity | MF |
| GO:0051087 | 0.03540539 | 1 | 2 | chaperone binding | MF |

| | | | | | |
|---|---|---|---|---|---|
| GO:0042626 | 0.04028636 | 2 | 20 | ATPase-coupled transmembrane transporter activity | MF |
| GO:0009055 | 0.04328176 | 4 | 78 | electron transfer activity | MF |
| GO:0003985 | 0.04942691 | 1 | 3 | acetyl-CoA C-acetyltransferase activity | MF |
| GO:0070573 | 0.04976681 | 1 | 3 | metallodipeptidase activity | MF |

**Supplementary Table S2**. List of underexpressed Gene Ontology functions

| category | pvalue | numDEInCat | numInCat | term | ontology |
|---|---|---|---|---|---|
| GO:0006412 | 8.84E-09 | 50 | 64 | translation | BP |
| GO:0007165 | 0.000510752 | 28 | 36 | signal transduction | BP |
| GO:0006782 | 0.000699073 | 10 | 10 | protoporphyrinogen IX biosynthetic process | BP |
| GO:0006096 | 0.000889921 | 10 | 10 | glycolytic process | BP |
| GO:0017004 | 0.003467099 | 10 | 11 | cytochrome complex assembly | BP |
| GO:0006935 | 0.003629206 | 29 | 41 | chemotaxis | BP |
| GO:0006417 | 0.004975633 | 6 | 6 | regulation of translation | BP |
| GO:0044205 | 0.006529388 | 7 | 7 | 'de novo' UMP biosynthetic process | BP |
| GO:0008615 | 0.012956241 | 6 | 6 | pyridoxine biosynthetic process | BP |
| GO:0006457 | 0.022597303 | 15 | 21 | protein folding | BP |
| GO:0042773 | 0.026715406 | 5 | 5 | ATP synthesis coupled electron transport | BP |
| GO:0006094 | 0.033515037 | 7 | 8 | gluconeogenesis | BP |
| GO:0045892 | 0.035586363 | 11 | 15 | negative regulation of transcription, DNA-templated | BP |
| GO:0006520 | 0.040308198 | 10 | 13 | cellular amino acid metabolic process | BP |
| GO:0006352 | 0.048860879 | 14 | 21 | DNA-templated transcription, initiation | BP |
| GO:0005840 | 2.71E-07 | 34 | 42 | ribosome | CC |
| GO:0005737 | 0.003253104 | 197 | 359 | cytoplasm | CC |
| GO:0032993 | 0.007105637 | 13 | 16 | protein-DNA complex | CC |
| GO:0015935 | 0.007811049 | 6 | 6 | small ribosomal subunit | CC |
| GO:0015934 | 0.013011445 | 7 | 8 | large ribosomal subunit | CC |

| | | | | | |
|---|---|---|---|---|---|
| GO:0045261 | 0.024393118 | 5 | 5 | proton-transporting ATP synthase complex, catalytic core F(1) | CC |
| GO:0019867 | 0.027048664 | 12 | 16 | outer membrane | CC |
| GO:0045263 | 0.037238775 | 4 | 4 | proton-transporting ATP synthase complex, coupling factor F(o) | CC |
| GO:0043190 | 0.048200913 | 34 | 56 | ATP-binding cassette (ABC) transporter complex | CC |
| GO:0003735 | 5.05E-10 | 45 | 54 | structural constituent of ribosome | MF |
| GO:0003700 | 1.11E-08 | 163 | 252 | DNA-binding transcription factor activity | MF |
| GO:0019843 | 5.93E-07 | 31 | 37 | rRNA binding | MF |
| GO:0000049 | 0.001342403 | 21 | 27 | tRNA binding | MF |
| GO:0000976 | 0.002366267 | 13 | 15 | transcription cis-regulatory region binding | MF |
| GO:0051082 | 0.002373321 | 11 | 12 | unfolded protein binding | MF |
| GO:0001217 | 0.007187085 | 9 | 10 | DNA-binding transcription repressor activity | MF |
| GO:0008137 | 0.008806499 | 9 | 10 | NADH dehydrogenase (ubiquinone) activity | MF |
| GO:0016987 | 0.009369077 | 18 | 25 | sigma factor activity | MF |
| GO:0046933 | 0.011855486 | 8 | 9 | proton-transporting ATP synthase activity, rotational mechanism | MF |
| GO:0048038 | 0.01598414 | 11 | 14 | quinone binding | MF |
| GO:0004519 | 0.031082771 | 11 | 15 | endonuclease activity | MF |
| GO:0004356 | 0.032923826 | 7 | 8 | glutamate-ammonia ligase activity | MF |
| GO:0020037 | 0.038913599 | 32 | 52 | heme binding | MF |
| GO:0008289 | 0.039735835 | 4 | 4 | lipid binding | MF |
| GO:0001216 | 0.041645693 | 8 | 10 | DNA-binding transcription activator activity | MF |
| GO:0050661 | 0.043662632 | 17 | 25 | NADP binding | MF |
| GO:0000287 | 0.044190909 | 45 | 76 | magnesium ion binding | MF |

**Supplementary Table S3**. List of primers used for the construction of the plasmids for the complementation experiments.

| Primer | Sequence | Gene | Restriction enzyme |
|---|---|---|---|
| EcoRI-PP_3998F | TGCCGGAATTCTTGTTTCAGGAGCTCACAA | *gqr* | EcoRI |
| HindIII-PP_3998R | CCCAAGCTTTCAAGCCTGCCAAATC | | HindIII |
| EcoRI-PP_4936F | TGCCGGAATTCATGTTGTATCAAAAAAACTGGGCA | *wzy* | EcoRI |
| HindIII-PP_4936R | CCCAAGCTTTCAGGTTGAACCGGCAATT | | HindIII |
| EcoRI-PP_4322F | TGCCGGAATTCTCATGCAGCTGCTCCA | *ccmF* | EcoRI |
| HindIII-PP_4322R | CCCAAGCTTTCATGCAGCTGCTCCA | | HindIII |
| BamHI-PP_4189F | TTCGCGGATCCATGCAAGAAAGCGTGATG | *sucA* | BamHI |
| HindIII-PP_4189R | CCCAAGCTTTTAGACAGTGAAGGCGTCTT | | HindIII |
| BamHI-PP_4935F | TTCGCGGATCCATGGCCGAAACACCG | *msbA* | BamHI |
| HindIII-PP_4935R | CCCAAGCTTTCAGGTGATATCGGCCTTG | | HindIII |
| EcoRI-PP_4799F | TGCCGGAATTCATGTACTGCGCCAAAACCTTC | *ldcA* | EcoRI |
| HindIII-PP_4799R | CCCAAGCTTTCACGCCCTGAACAGATC | | HindIII |
| EcoRI-PP_3999F | TGCCGGAATTCATGGATTACCTGCCGCTGT | *cysG* | EcoRI |
| HindIII-PP_3999R | CCCAAGCTTTCAGCTGTTCTGCGAACCTT | | HindIII |
| EcoRI-PP_5154 | TGCCGGAATTCATGACCCTTGTCGACCC | D2HGDH | EcoRI |
| HindIII-PP_5154R | CCCAAGCTTTTATTCAGGTGCGAAGATCT | | HindIII |

### Supplementary Figures

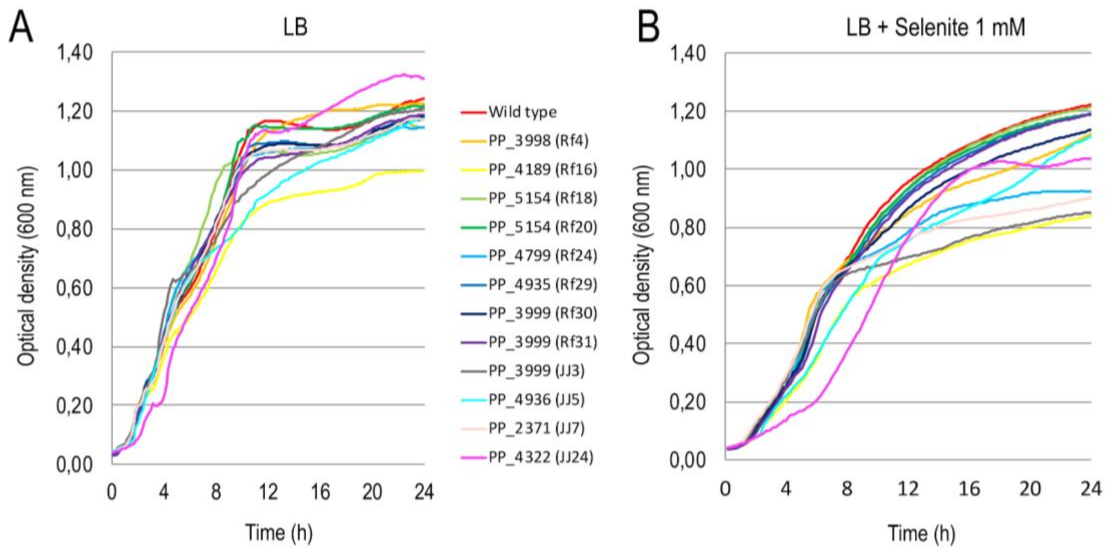


**Supplementary Figure S1. Growth curves of *P. putida* KT2440 and deficient strains in A) LB medium and B) LB medium supplemented with 1 mM selenite**. Bacterial growth was estimated on monitoring the optical density at 600nm (Synergy H1 Hybrid Multi-ModeReader, Biotek, Winooski VT, USA) over three biological replicates of cultures in plates (96 wells; Nunclon Δ Surface;Nunc A/S, Roskilde, Denmark). Plates were incubated at 30°C for 24h with continuous orbital shaking at 180 r.p.m.; the optical density was measured every 10min.

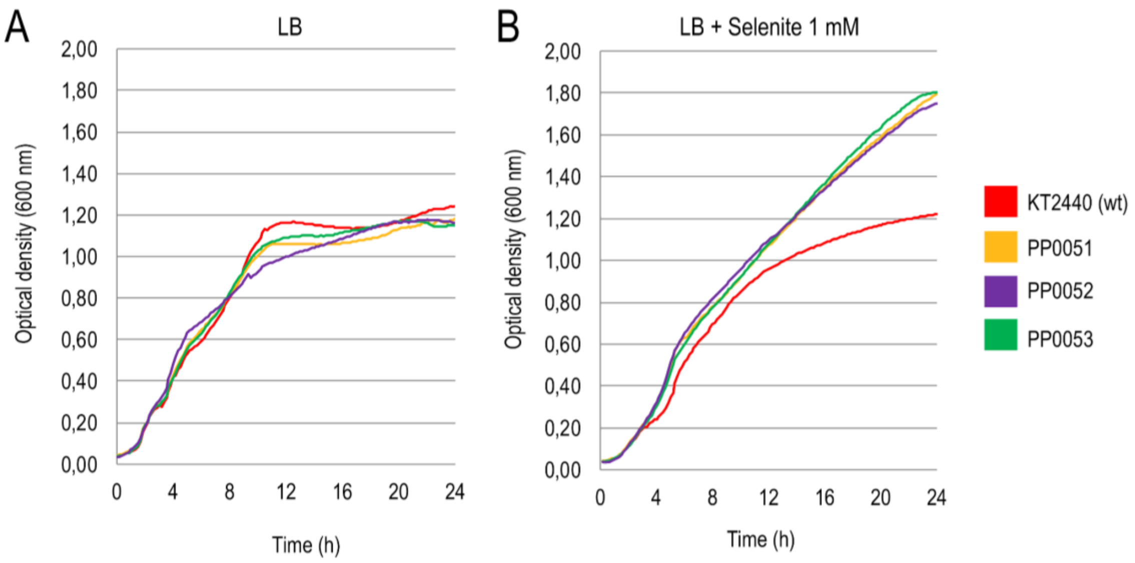


**Supplementary Figure S2. Growth curves of *P. putida* KT2440 and fast strains in A) LB medium and B) LB medium supplemented with 1 mM selenite**. Bacterial growth was estimated on monitoring the optical density at 600nm (Synergy H1 Hybrid Multi-ModeReader, Biotek, Winooski VT, USA) over three biological replicates of cultures in plates (96 wells; Nunclon Δ Surface;Nunc A/S, Roskilde, Denmark). Plates were incubated at 30°C for 24h with continuous orbital shaking at 180 r.p.m.; the optical density was measured every 10min.

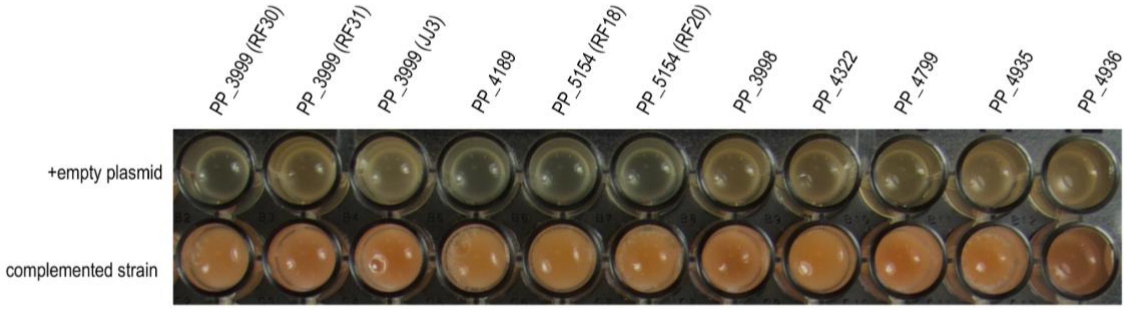


**Supplementary Figure S3. Growth of deficient strains complemented with the corresponding gene in trans in the presence of selenite**. The complementation experiments implied the expression of each gene with 3MB as inductor in the corresponding mutant strain (second row). Deficient strains carrying the pSEVA438 vector served as controls (first row). The photographs were taken after 24 h of culture in LB medium in the presence of selenite 1 mM and Sm 100 μg $ml^{-1}$.

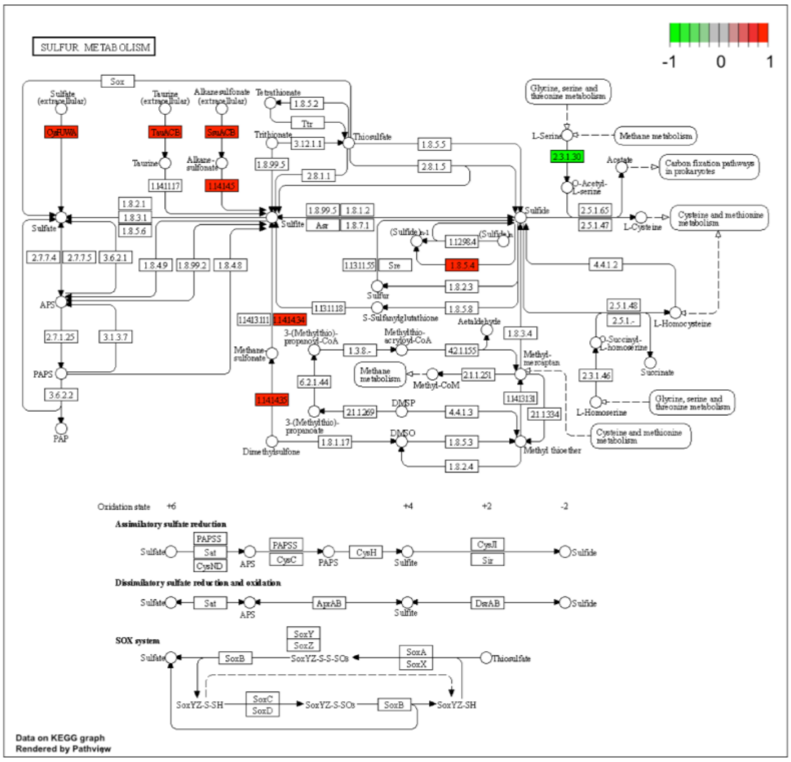


**Supplementary Figure S4. Reference pathways for sulfur metabolism obtained from The KEGG PATHWAY database**. Genes underexpressed in *P. putida* KT2440 in the presence of selenite in transcriptomic experiments are marked in green. Genes overexpressed in *P. putida* KT2440 in the presence of selenite in transcriptomic experiments are marked in red.

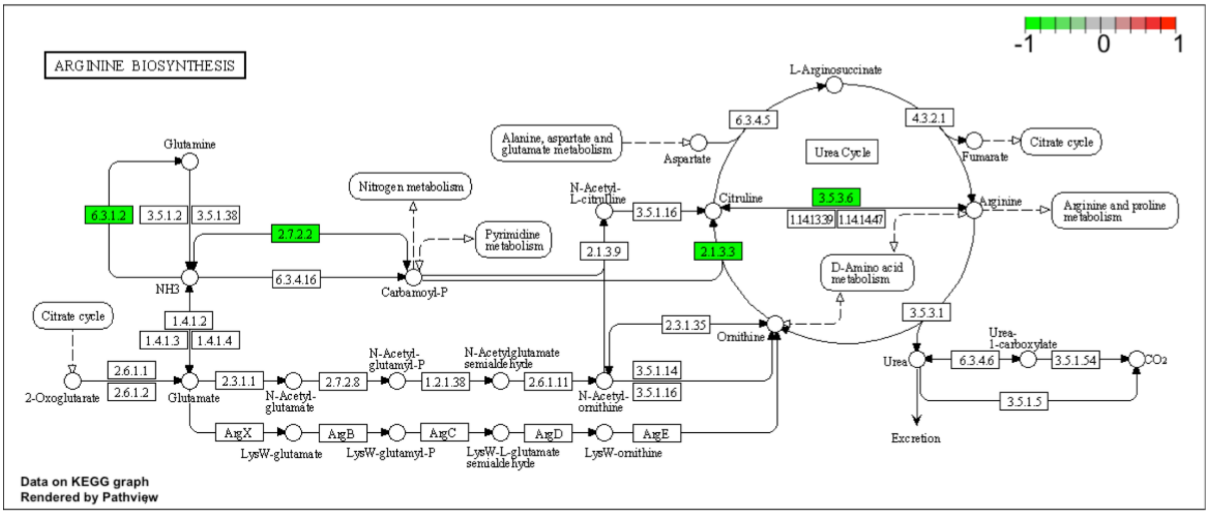


**Supplementary Figure S5. Reference pathways for arginine biosynthesis obtained from The KEGG PATHWAY database**. Genes underexpressed in *P. putida* KT2440 in the presence of selenite in transcriptomic experiments are marked in green.

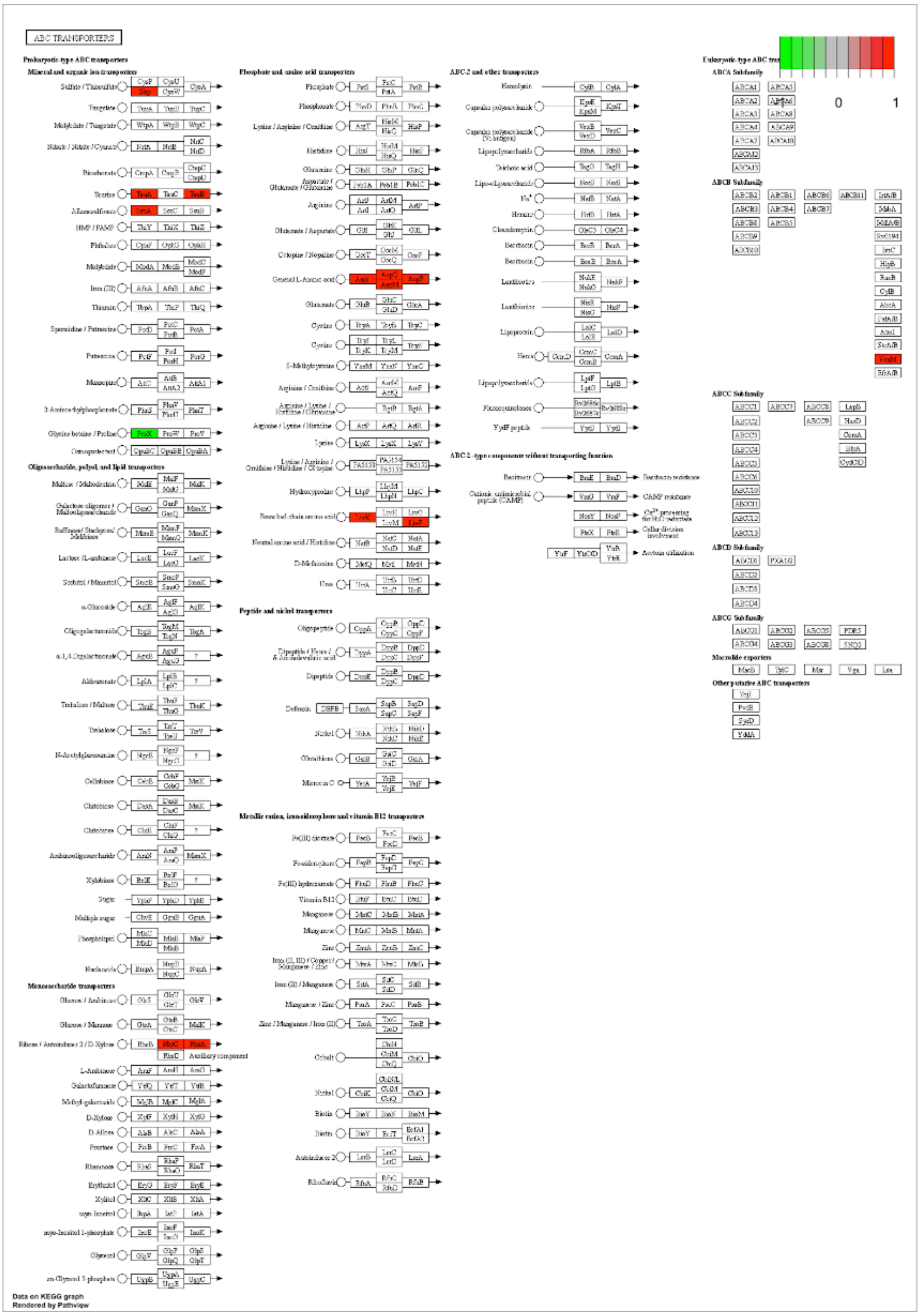


**Supplementary Figure S6. Reference for ABC transporters obtained from The KEGG PATHWAY database**. Genes underexpressed in *P. putida* KT2440 in the presence of selenite in transcriptomic experiments are marked in green. Genes overexpressed in *P. putida* KT2440 in the presence of selenite in transcriptomic experiments are marked in red.

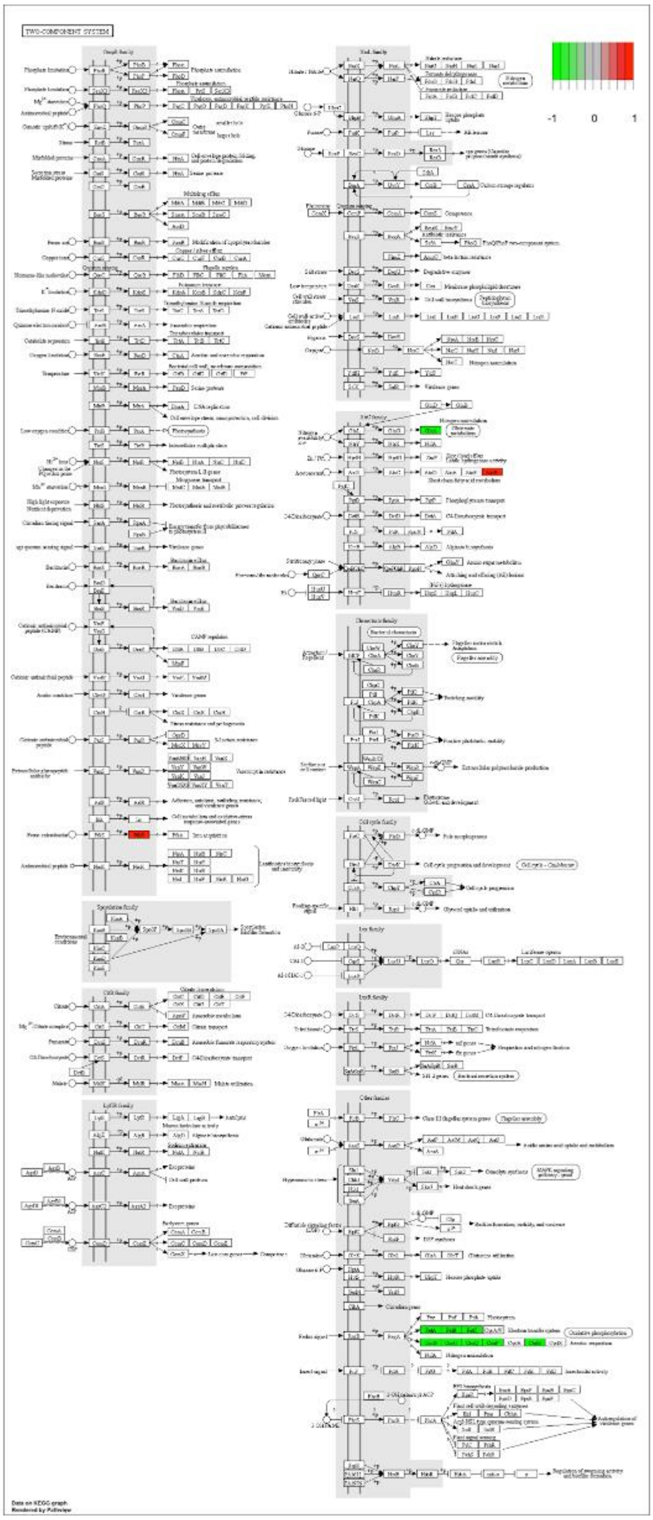


**Supplementary Figure S7. Reference for two-component systems obtained from The KEGG PATHWAY database**. Genes underexpressed in *P. putida* KT2440 in the presence of selenite in transcriptomic experiments are marked in green. Genes overexpressed in *P. putida* KT2440 in the presence of selenite in transcriptomic experiments are marked in red.

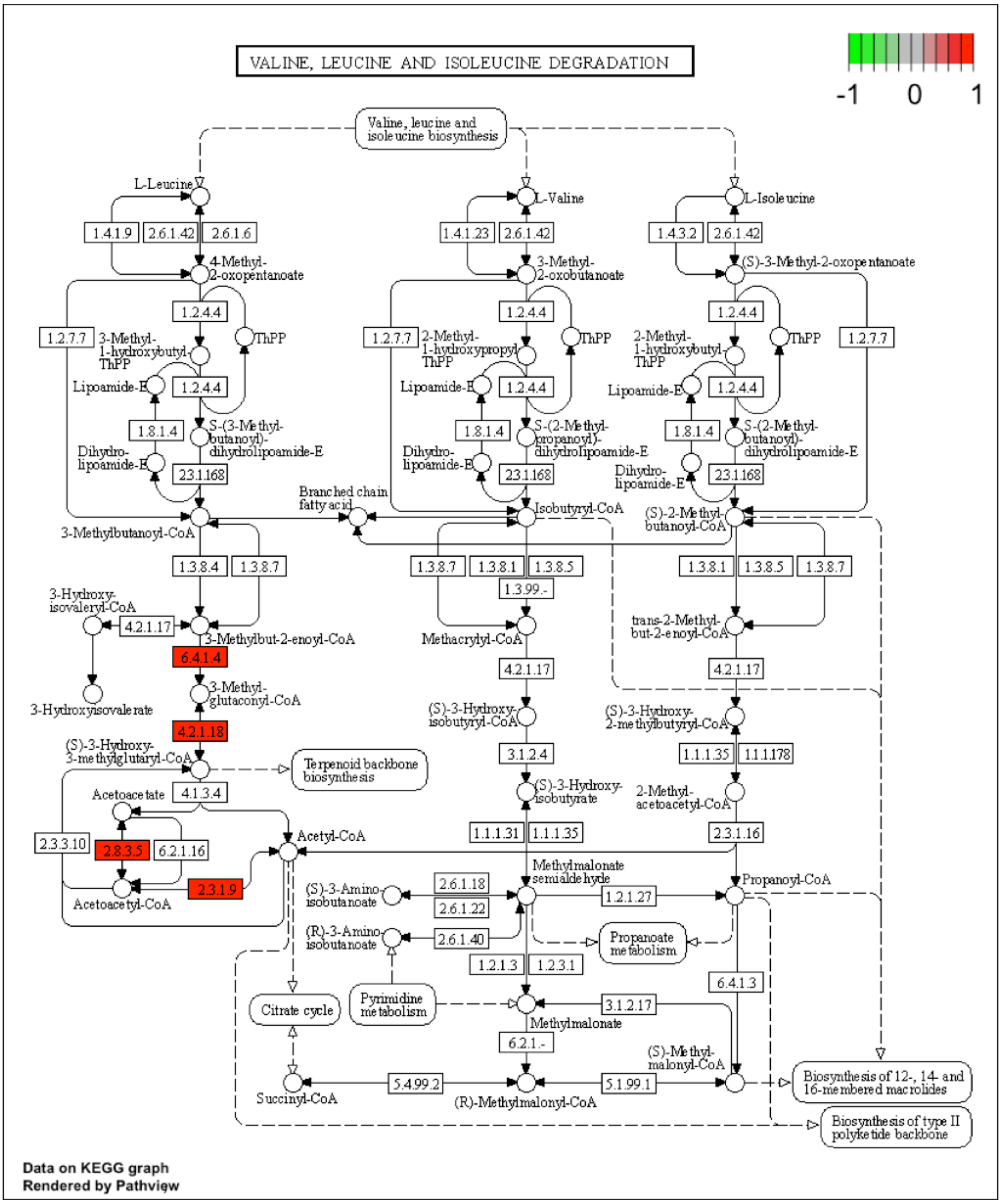


**Supplementary Figure S8**. **Reference pathways for valine, leucine and isoleucine degradation obtained from The KEGG PATHWAY database**. Genes overexpressed in *P. putida* KT2440 in the presence of selenite in transcriptomic experiments are marked in red.

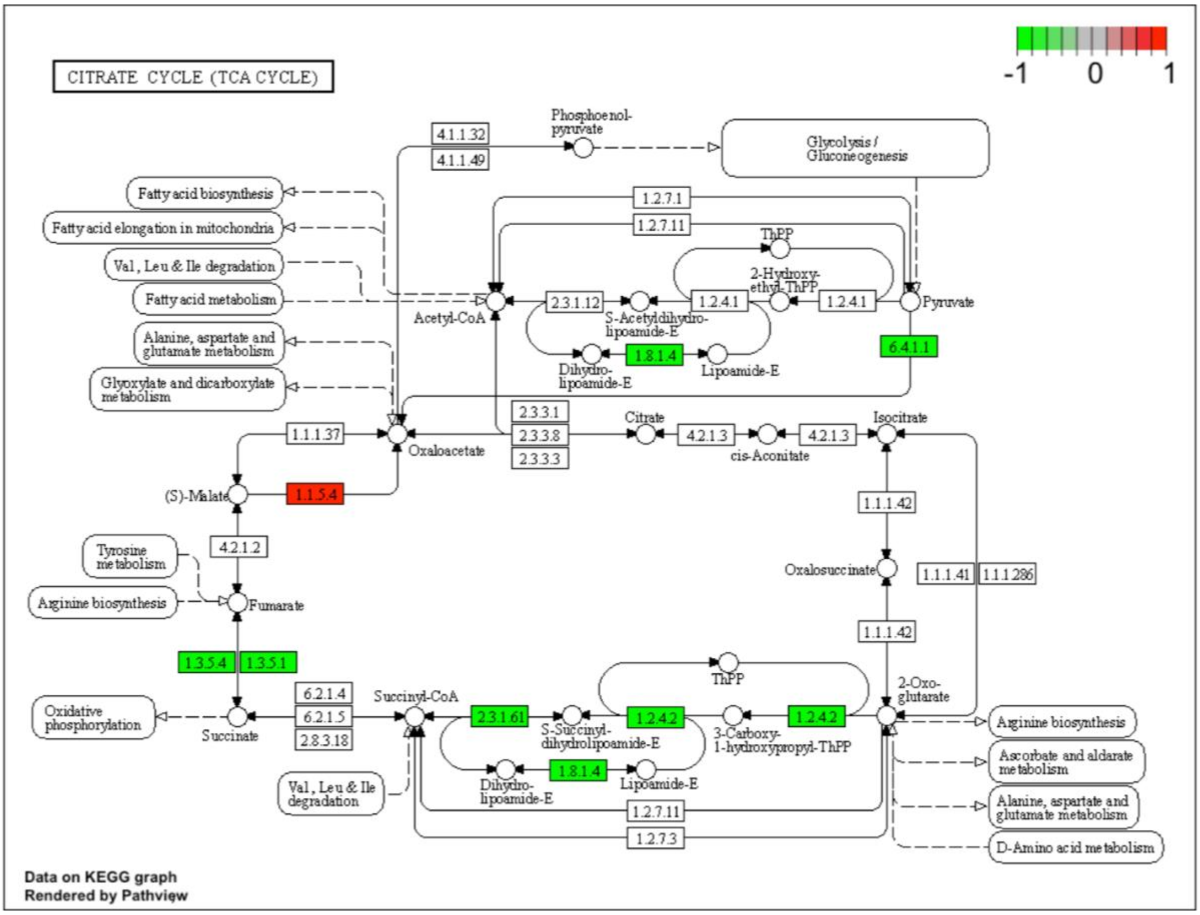


**Supplementary Figure S9. Reference pathway for TCA cycle obtained from The KEGG PATHWAY database**. Genes underexpressed in *P. putida* KT2440 in the presence of selenite in transcriptomic experiments are marked in green. Genes overexpressed in *P. putida* KT2440 in the presence of selenite in transcriptomic experiments are marked in red.

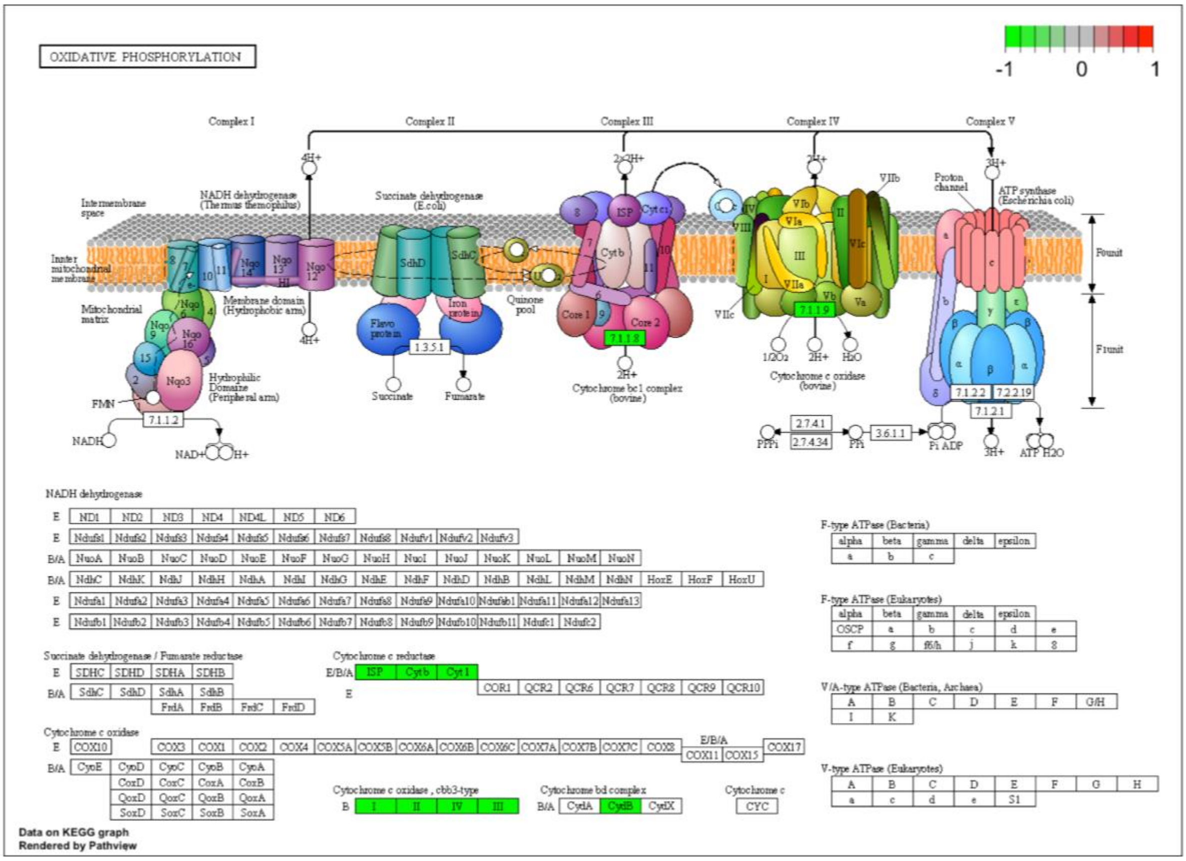


**Supplementary Figure S10. Reference pathways for oxidative phosphorylation obtained from The KEGG PATHWAY database**. Genes underexpressed in *P. putida* KT2440 in the presence of selenite in transcriptomic experiments are marked in green.

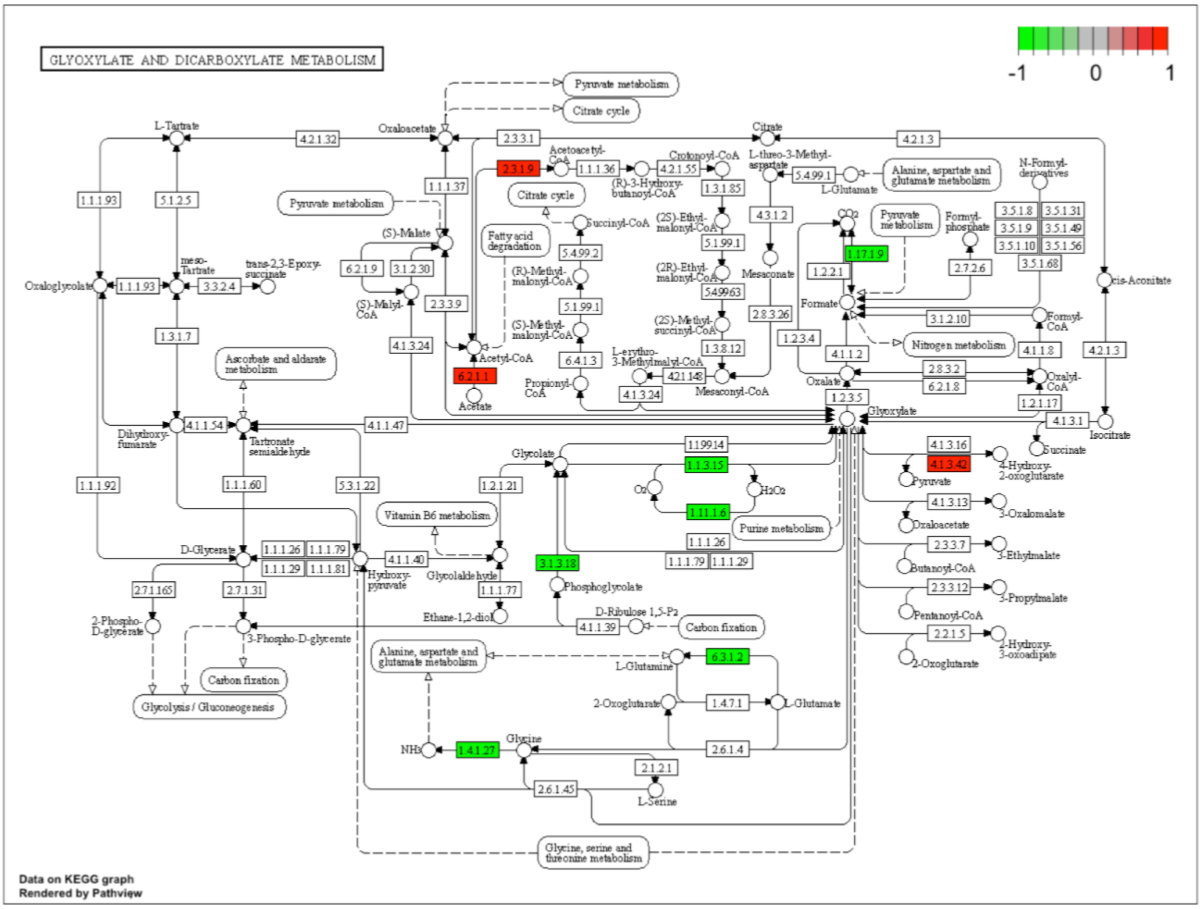


**Supplementary Figure S11. Reference pathways for glyoxylate and dicarboxylate obtained from The KEGG PATHWAY database**. Genes underexpressed in *P. putida* KT2440 in the presence of selenite in transcriptomic experiments are marked in green. Genes overexpressed in *P. putida* KT2440 in the presence of selenite in transcriptomic experiments are marked in red.

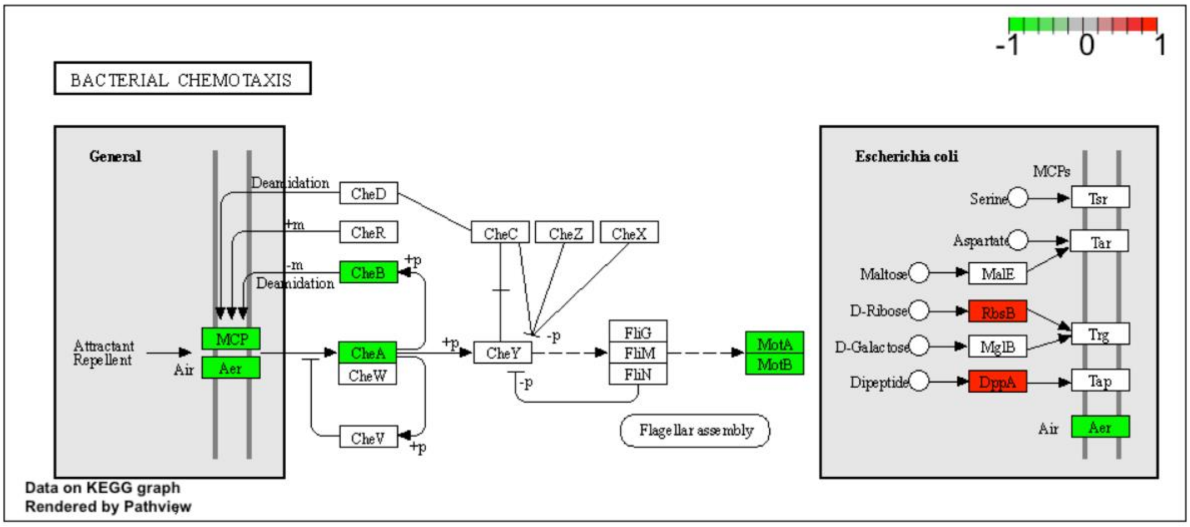


**Supplementary Figure S12. Reference pathways for bacterial chemotaxis obtained from The KEGG PATHWAY database**. Genes underexpressed in *P. putida* KT2440 in the presence of selenite in transcriptomic experiments are marked in green. Genes overexpressed in *P. putida* KT2440 in the presence of selenite in transcriptomic experiments are marked in red.

.
